# mTOR activation induces endolysosomal remodeling and nonclassical secretion of IL-32 via exosomes in inflammatory reactive astrocytes

**DOI:** 10.1101/2021.09.11.459904

**Authors:** Kun Leng, Brendan Rooney, Frank McCarthy, Wenlong Xia, Indigo V.L. Rose, Sophie Bax, Marcus Chin, Saeed Fathi, Kari A. Herrington, Manuel Leonetti, Aimee Kao, Stephen P.J. Fancy, Joshua E. Elias, Martin Kampmann

## Abstract

Astrocytes respond and contribute to neuroinflammation by adopting inflammatory reactive states. Although recent efforts have characterized the gene expression signatures associated with these reactive states, the cell biology underlying inflammatory reactive astrocyte phenotypes remains under-explored. Here, we used CRISPR-based screening in human iPSC-derived astrocytes to identify mTOR activation a driver of cytokine-induced endolysosomal system remodeling, manifesting as alkalinization of endolysosomal compartments, decreased autophagic flux, and increased exocytosis of certain endolysosomal cargos. Through endolysosomal proteomics, we identified and focused on one such cargo – IL-32, a disease-associated pro-inflammatory cytokine not present in rodents, whose secretion mechanism is not well understood. We found that IL-32 was partially secreted in extracellular vesicles likely to be exosomes. Furthermore, we found that IL-32 was involved in the polarization of inflammatory reactive astrocyte states, was upregulated in astrocytes in multiple sclerosis lesions, and preferentially co-localized with astrocytes in hypoxic-ischemic encephalopathy. We believe that our results advance our understanding of cell biological pathways underlying inflammatory reactive astrocyte phenotypes and identify potential therapeutic targets.

## INTRODUCTION

Astrocytes maintain homeostasis of the central nervous system in myriad ways, for example through phagocytosis of synapses (1,2) or exocytosis of ATP or glutamate (3–5). In the context of central nervous system injury or disease, astrocytes respond to and amplify inflammatory signaling cascades (6,7), adopting inflammatory reactive astrocyte states characterized by distinct gene expression and cytokine signatures (8–10). Although recent efforts have elucidated these signatures, the cell biological pathways underlying inflammatory reactive astrocyte phenotypes remain under-explored.

The endolysosomal system encompasses a diverse pool of intracellular vesicles of varying luminal pH that mediate degradative (e.g. lysosomes, phagosomes, autophagosomes) as well as exocytic functions (e.g. recycling endosomes, multivesicular bodies) (11,12). Although a large body of work exists on the cross-regulation of autophagy and inflammatory responses in immune cells (13), much less is known about how this occurs in brain cell types, especially astrocytes.

Here, using human induced pluripotent stem cell (hiPSC)-derived astrocytes, we found that the inflammatory cytokines IL-1α+TNF+C1q in combination (hereafter abbreviated as “ITC”), which have been widely used to induce inflammatory astrocyte reactivity in vitro (14), caused mTOR activation and mTOR-dependent remodeling of the endolysosomal system, which was associated with increased exocytosis of certain endolysosomal cargos. Through endolysosomal proteomics, we identified and focused on one such cargo, the pro-inflammatory cytokine IL-32. Although cerebrospinal fluid levels of IL-32 are elevated in neuroinflammatory diseases such as multiple sclerosis (15), the secretion mechanism of IL-32 is not well understood, as it lacks a classical signal peptide (16,17). Furthermore, it is unclear what cell types in the central nervous system produce IL-32 under neuroinflammatory conditions. We found that IL-32 was upregulated in astrocytes in multiple sclerosis lesions, was likely secreted in part via exosomes, and was involved in the polarization of inflammatory reactive astrocyte states.

## RESULTS

To identify the cellular pathways responsible for the loss of homeostatic functions that accompany inflammatory astrocyte reactivity, we reanalyzed RNA-sequencing data from multiple hiPSC-derived astrocyte models (18–22) (Supplementary Table 1) treated with ITC or similar treatments (Fig. 1a), focusing on downregulated genes. We found that downregulated genes after ITC treatment were enriched in genes encoding proteins associated with the endolysosomal system (Fig. 1b-c, Supplementary Fig. 1), suggesting a potential link to the reported phenotype of decreased phagocytosis.

**Figure 1.**
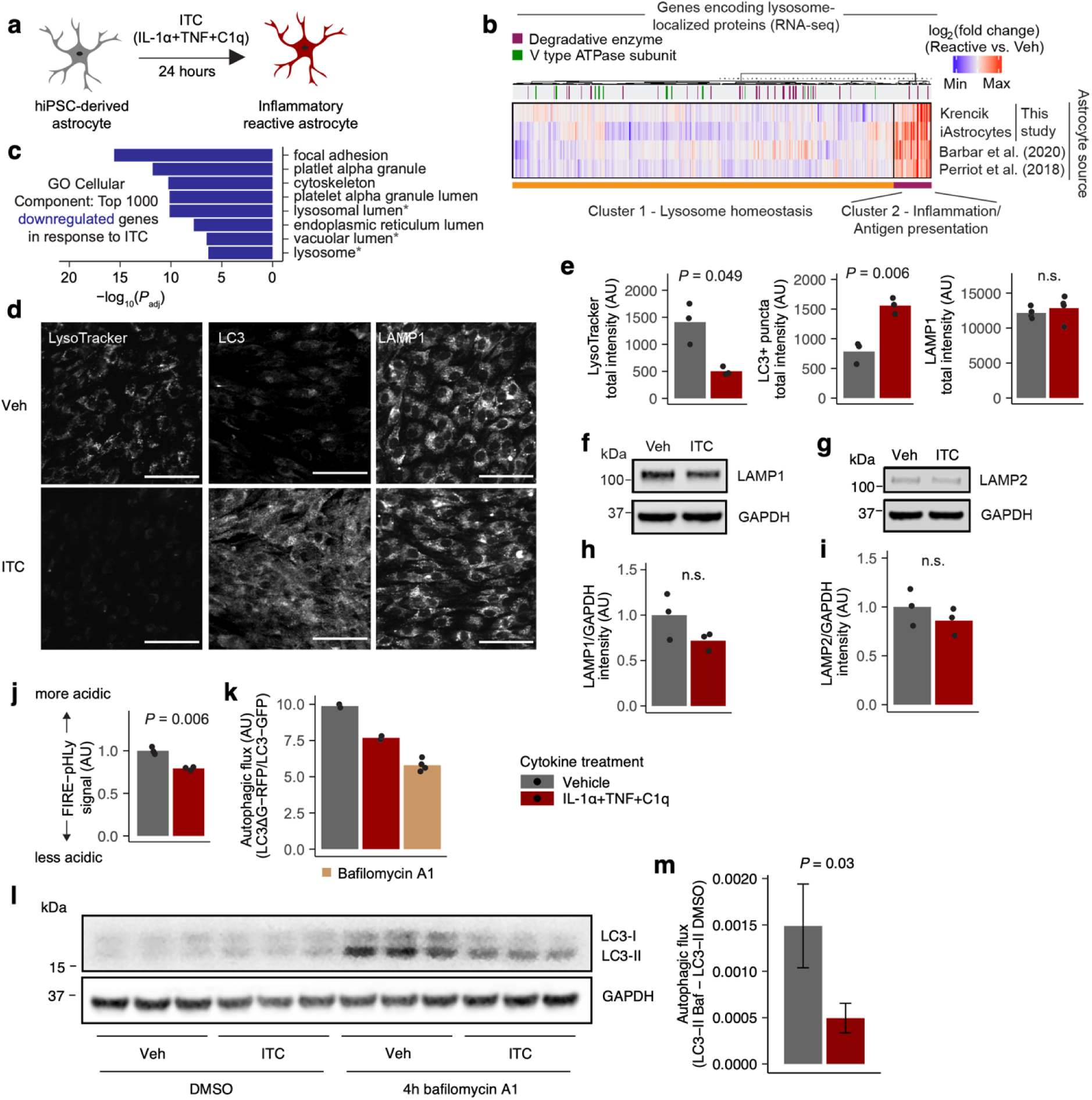
Endolysosomal function is perturbed in inflammatory reactive astrocytes. **a**, Schematic of modeling inflammatory astrocyte reactivity in vitro. **b**, Heatmap of changes in the expression of genes encoding lysosome-localized proteins in different hiPSC-derived astrocyte models treated with ITC or similar treatments vs vehicle control. **c**, Top GO Cellular Component terms enriched among the top 1000 downregulated genes in iAstrocytes treated with ITC; endolysosomal system-related terms are highlighted by asterisks. **d**, Representative images of LysoTracker staining in live iAstrocytes or immunostaining of LC3 or LAMP1 in fixed and permeabilized iAstrocytes; scale bar = 75 um. **e**, Quantification of imaging experiments shown in **d** (n = 3 wells per condition). **f**-**g**, Representative immunoblot bands against LAMP1 (**f**) or LAMP2 (**g**). **h**-**i**, Quantification of immunoblot experiments shown in **f**-**g** (n = 3 wells per condition). **j**, Measurement of the acidity of LAMP1^+^ endolysosomal compartments using FIRE-pHLy via flow cytometry (see Methods). **k**, Measurement of autophagic flux with the LC3ΔG-RFP/LC3-GFP fluorescent reporter from Kaizuka *et al*. (25). **l**, Immunoblot against LC3 demonstrating LC3-I and LC3-II bands. **m**, Quantification of autophagic flux from the LC3-II bands in **l**; error bars reflect the 95% confidence interval associated with the standard error of the mean; individual data points not shown because the quantities of interest are differences between means, with no biologically meaningful pairing of individual data points across conditions. *P* values where shown were calculated using two-sided Student’s t test.

To confirm if ITC indeed perturbed endolysosomal system function, we performed downstream experiments using the iAstrocyte model that we previously developed (WTC11 genotype) (9). We found that ITC reduced LysoTracker staining without appreciably changing total endolysosomal mass as measured by the abundance of LAMP1 or LAMP2 (Fig. 1d-i, Supplementary File 1), which tend to be enriched in lysosomes but can also mark other endolysosomal compartments (23). Since a decrease in LysoTracker staining can reflect a decrease in either total endolysosomal mass or acidity, we deduced from the above results that ITC caused endolysosomal alkalinization. Indeed, using a genetically encoded endolysosomal pH reporter (FIRE-pHLy) (24), we found that ITC decreased the acidity of LAMP1^+^ endolysosomal compartments (Fig. 1j). We also observed a buildup of LC3^+^ puncta on immunostaining after ITC (Fig. 1d-e), suggestive of perturbed autophagic flux. Using both a genetically encoded reporter (25) (Fig. 1k) and LC3-II western blot (26) (Fig. 1l-m, Supplementary File 1), we found that ITC decreased autophagic flux as well, consistent with the importance of acidic lysosomes for degradation of autophagic substrates (27).

To gain a more detailed understanding of how endolysosomal function is perturbed in inflammatory reactive astrocytes, we performed mass spectrometry-based proteomic characterization of LAMP1^+^ endolysosomal compartments in ITC-treated vs control iAstrocytes (Fig. 2a, see Methods), using the total cell lysate as a reference. We found that v-ATPase subunits and degradative enzymes were less abundant in endolysosomal compartments in ITC-treated iAstrocytes (Fig. 2b-c), consistent with transcript-level downregulation in the RNA-seq data (Fig. 1b). Furthermore, CST3, a potent inhibitor of lysosomal proteases (28), accumulated in endolysosomal compartments upon ITC treatment (Supplementary Table 2). We also detected endolysosomal accumulation of proteins involved in vesicular exocytosis (e.g. RAB27A (29), SNAP25 (3,5)) together with inflammatory mediators (e.g. IL-32 (16,17), CCL2 (30)), suggesting that ITC may cause rerouting of endolysosomal trafficking to facilitate nonclassical secretion of inflammatory factors.

**Figure 2.**
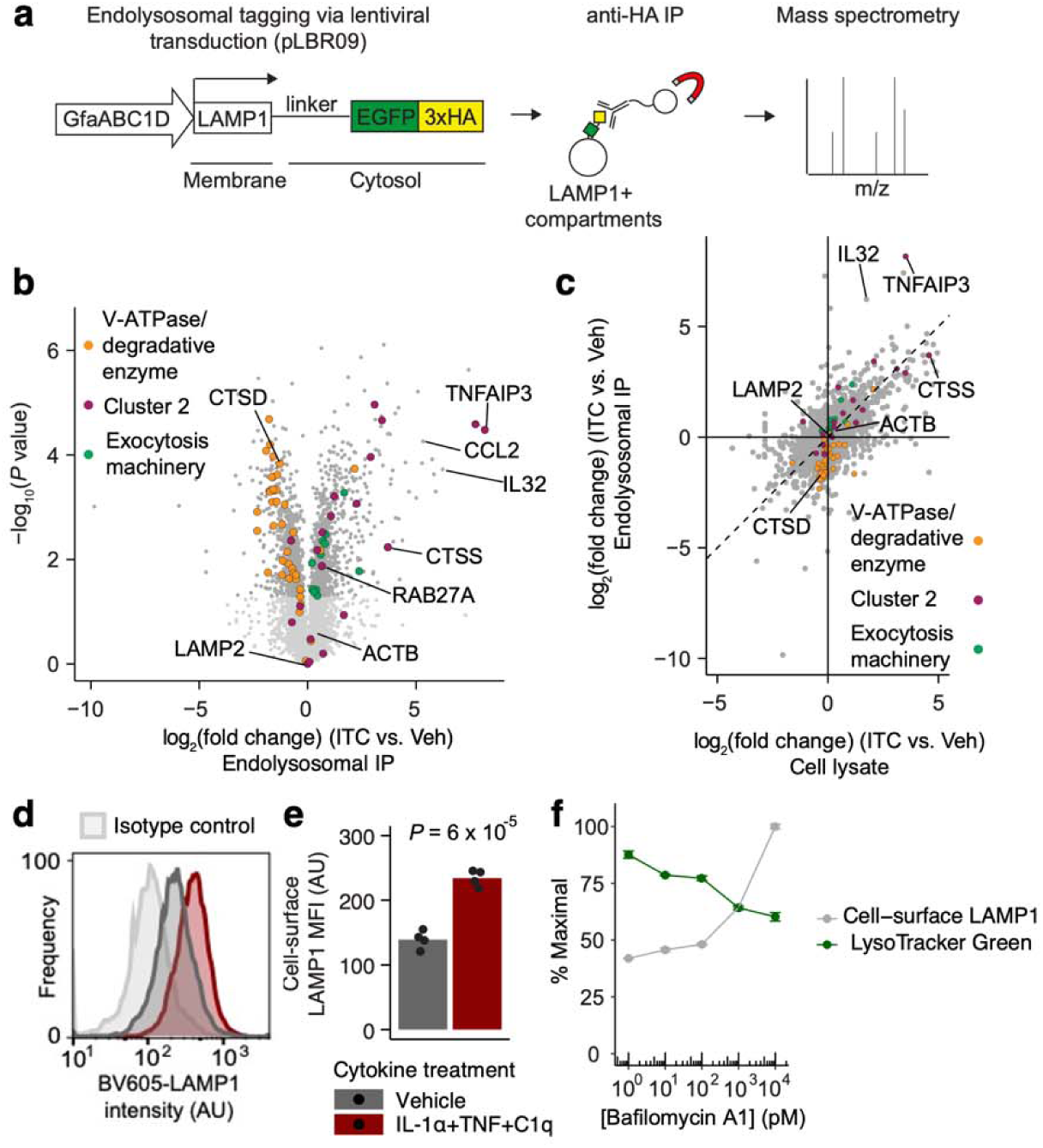
Perturbation of endolysosomal function is accompanied by remodeling of the endolysosomal proteome. **a**, Schematic of endolysosomal proteomic workflow (n = 3 wells per condition). **b**, Volcano plot of endolysosomal proteomic data. **c**, Scatterplot comparing log_2_-fold change of endolysosomal vs total cell lysate protein abundance in ITC-treated iAstrocytes compared to vehicle-treated iAstrocytes (IP – immunoprecipitation). **d**, Representative histograms of cell-surface LAMP1 staining intensity in vehicle- or ITC-treated iAstrocytes compared to isotype control staining intensity measured by flow cytometry. **e**, Median fluorescence intensity (MFI) of cell-surface LAMP1 measured by flow cytometry (n = 3 wells per condition); *P* value by two-sided Student’s t test. **f**, Cell-surface LAMP1 or LysoTracker staining in iAstrocytes treated with increasing doses of bafilomycin A1 (n = 3 wells per condition; error bars reflect the standard error of the mean).

To explore this hypothesis, we developed a flow-cytometric assay to quantify in live unpermeabilized iAstrocytes the exposure of the LAMP1 luminal domain on the plasma membrane (see Methods), which is known to increase in degranulating immune cells (31) and may be useful as a marker of the exocytic activity of certain endolysosomal pathways (32). With this approach, we found that ITC-treated iAstrocytes had ∼1.8 times the amount of cell-surface LAMP1 relative to vehicle-treated controls, despite equivalent levels of total LAMP1 (Figure 2d-e). We were able to directly observe putative endolysosomal exocytic events from LAMP1^+^ vesicles via total internal reflection fluorescence microscopy of iAstrocytes transduced with a LAMP1-mCherry fusion construct and loaded with LysoTracker Green, where the green fluorescence from LysoTracker was lost from LAMP1-mCherry^+^ vesicles trafficked to the plasma membrane (Supplementary File 2). Furthermore, the abundance of cell-surface LAMP1 demonstrated a positive dose-response relationship with bafilomycin A1 (Fig. 2f), an inhibitor of v-ATPase activity and autophagosome-lysosome fusion (33) that is known to increase endolysosomal exocytosis (34–36). Compared to cell-surface LAMP1, LysoTracker staining demonstrated a negative dose-response relationship (Fig. 2f), likely reflecting alkalinization of endolysosomal compartments due to v-ATPase inhibition.

To identify factors downstream of ITC that mediate the perturbations to endolysosomal function we observed above, we proceeded to perform targeted CRISPR-based inhibition (CRISPRi) screens against the “druggable genome” (37) in iAstrocytes treated with vehicle or ITC (see Methods) using cell-surface LAMP1 or LysoTracker staining as proxies of endolysosomal system function (Fig. 3a). Consistent with our observations from the bafilomycin A1 titration curve, the phenotype scores (labeled as “gene score”, see Methods) of hits from the cell-surface LAMP1 screens were on average inversely correlated with those from the LysoTracker screens (Fig. 3b,d). Notably, *MTOR* was a top hit in both screens (Fig. 3d), and we also found a strong enrichment for genes associated with the mTOR pathway in the top hits from both screens (Fig. 3c-d). These results corroborate in hiPSC-derived astrocytes the rich literature on the regulation of autophagy and lysosome function by mTOR (38,39), much of which was based on experiments in transformed human cell lines, yeast, or animal models.

**Figure 3.**
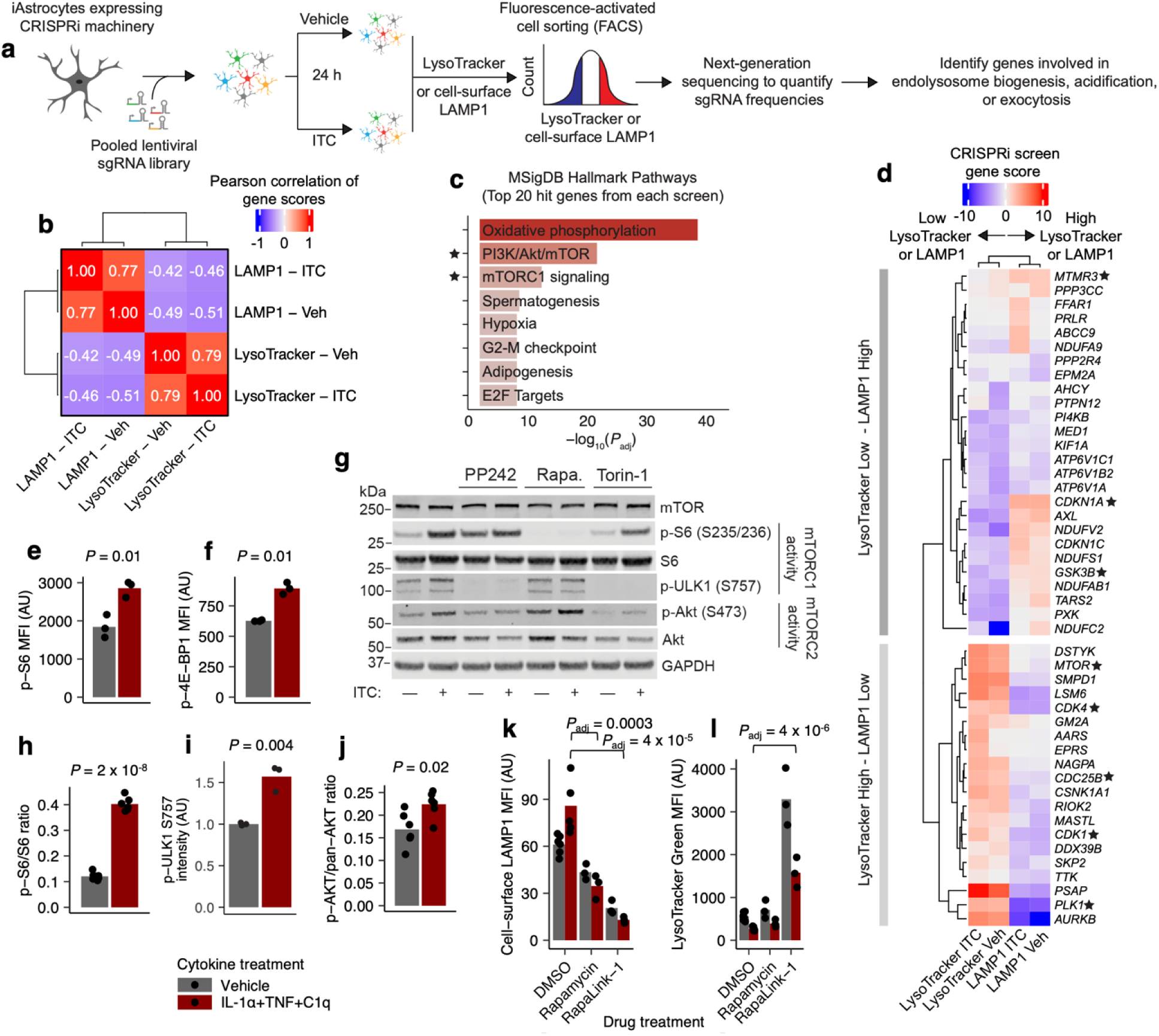
Multi-phenotypic CRISPR-based screening identifies mTOR as a central regulator of endolysosomal system function. **a**, Schematic of CRISPR-based screening workflow. **b**, Pearson correlation of gene scores (see Methods) of hits from the LAMP1 vs LysoTracker screens (n = 2 biological replicate screens per condition). **c**, Enrichment analysis against MSigDB Hallmark Pathways terms of the top 20 hits from each screen; terms pertaining to mTOR are highlighted with stars. **d**, Heatmap of gene scores of the hits overlapping with the highlighted MSigDB terms in **c**. **e**-**f**, Median fluorescence intensity (MFI) of phospho-S6 (**e**) or pospho-4E-BP1 staining in ITC-vs. vehicle-treated iAstrocytes measured by flow cytometry. **g**, Representative immunoblot bands corresponding to mTOR, phospho-S6, total S6, phospho-ULK1 and phospho-AKT1 in ITC- vs vehicle-treated iAstrocytes in the presence of different mTOR inhibitors. **h**, Quantification of immunoblotting experiments (n = 6 wells per condition for phospho-S6/S6, n = 3 wells per condition for phospho-ULK1, n = 6 wells per condition for pospho-AKT1/pan-AKT; *P* values from two-sided Student’s t test). **k**-**l**, Cell-surface LAMP1 (**k**) or LysoTracker (**l**) MFI in ITC- vs. vehicle-treated iAstrocytes in the presence of mTOR inhibitors measured by flow cytometry (n = 6 wells for DMSO treated, n = 3 wells for all other conditions; *P* values calculated only for ITC-treated conditions by linear regression with adjustment for multiple testing by Holm’s method, shown only if significant).

To validate our screening results, we first assessed whether ITC increased the activity of mTORC1or mTORC2, the two protein complexes formed by mTOR with distinct regulatory subunits and downstream signaling pathways (40). We found that ITC increased mTORC1 activity as measured by increased levels of Ser235/236-phosphoryated S6 ribosomal protein (hereon referred to as phospho-S6), Thr37/46-phosphorylated 4E-BP1, and Ser757-phosphorylated ULK1 (38,41) (Fig. 3e-i). ITC also increased mTORC2 activity as measured by Ser473-phosphorylated AKT1 (42) (Fig. 3j). To see if there was evidence of mTOR activation in inflammatory reactive astrocytes more broadly across different contexts, models, and species, we extracted differentially expressed genes in astrocytes from a variety of published rodent and human transcriptomic datasets and found broad enrichment for mTOR pathway-associated genes among upregulated genes (Supplementary Fig. 2), including in a recently published dataset demonstrating a role for astrocyte mTOR activation in experimental autoimmune encephalitis (EAE) (43), a mouse model of multiple sclerosis. Next, we tested the effect of mTOR inhibitors on cell-surface LAMP1 and LysoTracker staining. Consistent with the phenotype scores observed for *MTOR* knockdown in our CRISPRi screens, both rapamycin and Rapalink-1 (44) decreased cell-surface LAMP1 levels and increased LysoTracker staining (Fig. 3k-l), regardless of ITC treatment, although the effect of rapamycin on LysoTracker staining was not statistically significant. Given our factorial experimental design, we also analyzed our data with two-way ANOVA (see Methods). Rapamycin and Rapalink-1 decreased both baseline cell-surface LAMP1 (statistically significant main effect terms) as well as reversing the ITC-induced increase in cell-surface LAMP1 (statistically significant interaction terms) (Supplementary Table 4, tab Fig3k). For LysoTracker staining, Rapalink-1 increased both baseline LysoTracker staining as well as exacerbating the ITC-induced decrease in LysoTracker staining, but the effects of rapamycin were not statistically significant (Supplementary Table 4, tab Fig3l). Both drugs significantly decreased phospho-S6 levels at baseline as well as abrogating the ITC-induced increase (Supplementary Fig. 3a; Supplementary Table 4, tab SupFig3a), demonstrating target engagement.

Focusing on mTOR as a central regulator of endolysosomal system function, we next explored how modulating mTORC1 or mTORC2 activity affected the exocytic activity of endolysosomal pathways. First, we tested whether the changes in cell-surface LAMP1 and LysoTracker staining caused by ITC depended on mTORC1 vs. mTORC2. We found that knockdown of *MTOR* decreased cell-surface LAMP1 and increased LysoTracker staining in ITC-treated astrocytes, which was phenocopied to a larger degree by knockdown of *RPTOR* (which encodes a subunit unique mTORC1) (45) than by knockdown of *RICTOR* (which encodes a subunit unique to mTORC2) (45) (Fig. 4a), consistent with known mTORC1-dependent mechanisms regulating autophagy and lysosome function (38,39). We verified that we achieved robust knockdown of mTOR by both directly measuring mTOR protein levels as well as downstream phospho-S6 levels (Supplementary Fig. 3b-c). A robust decrease in downstream phospho-S6 levels with *RPTOR* knockdown also verified Raptor protein depletion.

**Figure 4.**
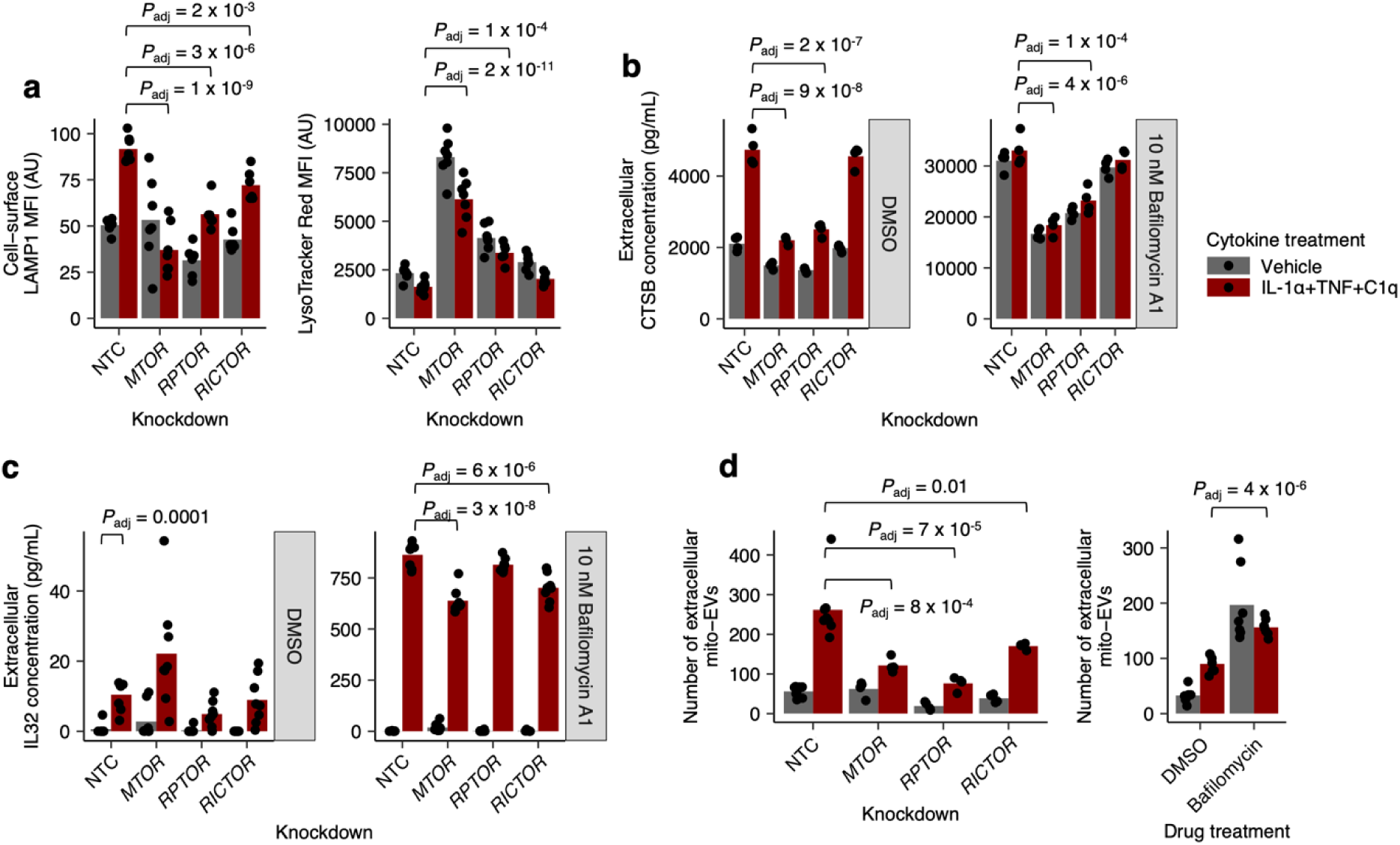
Cytokine-induced remodeling or pharmacological disruption of the endolyosomal system results in increased exocytosis of endolysosomal contents. **a**-**d**, Cell-surface LAMP1 or LysoTracker median fluorescence intensity (MFI) measured by flow cytometry (**a**), extracellular CTSB concentration measured by electrochemiluminescence-based immunoassay (**b**), extracellular IL-32 concentration measured by ELISA (**c**), or abundance of extracellular mito-EVs measured by flow cytometry (**d**) in ITC- vs. vehicle-treated iAstrocytes transduced with non-targeting control (NTC) sgRNAs or sgRNAs targeting genes encoding common (*MTOR*) or unique mTORC1 (*RPTOR*) vs mTORC2 (*RICTOR*) subunits, with or without co-treatment with bafilomycin A1. *P* values were calculated by linear regression with correction for multiple testing using Holm’s method, shown only when significant.

As we did previously with mTOR inhibitors, we also analyzed the mTORC1/2 subunit knockdown data by two-way ANOVA. Whereas the interaction between *MTOR* knockdown and ITC-treatment was statistically significant for both cell-surface LAMP1 and LysoTracker staining, the interactions between ITC treatment and *RPTOR* or *RICTOR* knockdown were not statistically significant for either cell-surface LAMP1 or LysoTracker staining (Supplementary Table 4, tabs Fig4a-b). This discrepancy suggests that knockdown of *RPTOR* or *RICTOR* may not cleanly inhibit mTORC1 or mTORC2 activity respectively without causing second-order changes. Indeed, we saw that *RICTOR* knockdown caused an increase in phospho-S6 levels and mTOR protein levels (Supplementary Fig. 3b-c), demonstrating that blocking mTORC2 activity can cause a compensatory increase in mTORC1 activity. Thus, phenotypes associated with *RPTOR* or *RICTOR* knockdown need to be interpreted with caution, as they likely do not reflect the effect of perturbing *only* mTORC1 or mTORC2 activity, respectively.

In addition to decreasing mTORC1 activity with *RPTOR* knockdown, we also tested the effect of increasing mTORC1 activity through knockdown of *TSC1*, an upstream inhibitor of mTORC1 (41). We found that *TSC1* knockdown increased cell-surface LAMP1 and decreased LysoTracker staining (Supplementary Fig. 4a-b), as one would predict. However, similar to what we observed with *RPTOR* knockdown, the interaction term between *TSC1* knockdown and ITC-treatment was not statistically significant (Supplementary Table 4, tabs SupFig4a-b).

Next, we assessed how mTORC1 vs. mTORC2 controlled the exocytic activity of endolysosomal pathways by measuring the extracellular levels of three different endolysosomal cargos (see Methods): cathepsin B (CTSB), a lysosome-enriched protease known to be released extracellularly by astrocytes (46); IL-32, which we identified as being enriched in endolysosomal compartments from our proteomics dataset, consistent with prior observations (47); and mitochondria-containing extracellular vesicles (mito-EVs) (see Methods and Supplementary Fig. 5), which are known to be released by astrocytes and are thought to be derived from mitophagy (48,49). We found that ITC increased extracellular levels of CTSB, IL-32, as well as mito-EVs, all of which were dramatically boosted by addition of bafilomycin A1 (Fig. 4b-d). Furthermore, *MTOR* knockdown decreased the exocytosis of CTSB, IL-32, and mito-EVs, although for IL-32 we only observed a statistically significant effect in the presence of bafilomycin A1 (Fig. 4b-d). *RPTOR* knockdown phenocopied *MTOR* knockdown to a larger degree than *RICTOR* knockdown for CTSB and mito-EVs, whereas for IL-32 *RICTOR* knockdown had a stronger effect than *RPTOR* knockdown (Fig. 4b-d). This trend was also evident by examining the interaction terms for *MTOR* vs *RICTOR* vs *RPTOR* knockdown with ITC treatment: for CTSB and mito-EVs, the interaction between ITC treatment and *MTOR* or *RPTOR* knockdown, but not *RICTOR* knockdown, was statistically significant (in the absence of bafilomycin A1); whereas for IL-32, the interaction ITC treatment and *RICTOR* knockdown, but not *MTOR* or *RPTOR* knockdown, was statistically significant (in the presence of bafilomycin A1) (Supplementary Table 4, tabs Fig4b-d). On the other hand, *TSC1* knockdown increased the exocytosis of mito-EVs and CTSB (Supplementary Fig. 4c-d), as expected, with statistically significant interaction terms as well (Supplementary Table 4, tabs SupFig4c-d). Overall, the stronger dependence on mTORC1 for the exocytosis of CTSB and mito-EVs compared to IL-32 suggests that IL-32 exocytosis may occur via a different endolysosomal pathway compared to CTSB and mito-EVs.

Given prior work which suggested that extracellular IL-32 is membrane-associated and possibly a component of extracellular vesicles (47,50,51), we proceeded to further characterize the secretion mechanism of IL-32 by isolating extracellular vesicles (EVs) from iAstrocyte conditioned media via differential ultracentrifugation (see Methods), with or without ITC treatment. By western blotting, we verified that our EV preparation contained high levels of consensus EV markers such as CD63, CD81, Hsc70, and caveolin-1 (52) and was of acceptable purity, containing undetectable levels of apo-lipoproteins such as ApoA-I (Fig. 5a, Supplementary File 1), in accordance with the MISEV guidelines (52). On nanoparticle tracking analysis (see Methods), we saw that the size distribution of EVs from our preparations centered around a diameter of 100 nm, as expected (52) (Fig. 5b). Both western blotting and nanoparticle tracking analysis suggested an increase in the concentration of EVs with ITC treatment (Fig. 5a-b, Supplementary Fig. 6).

**Figure 5.**
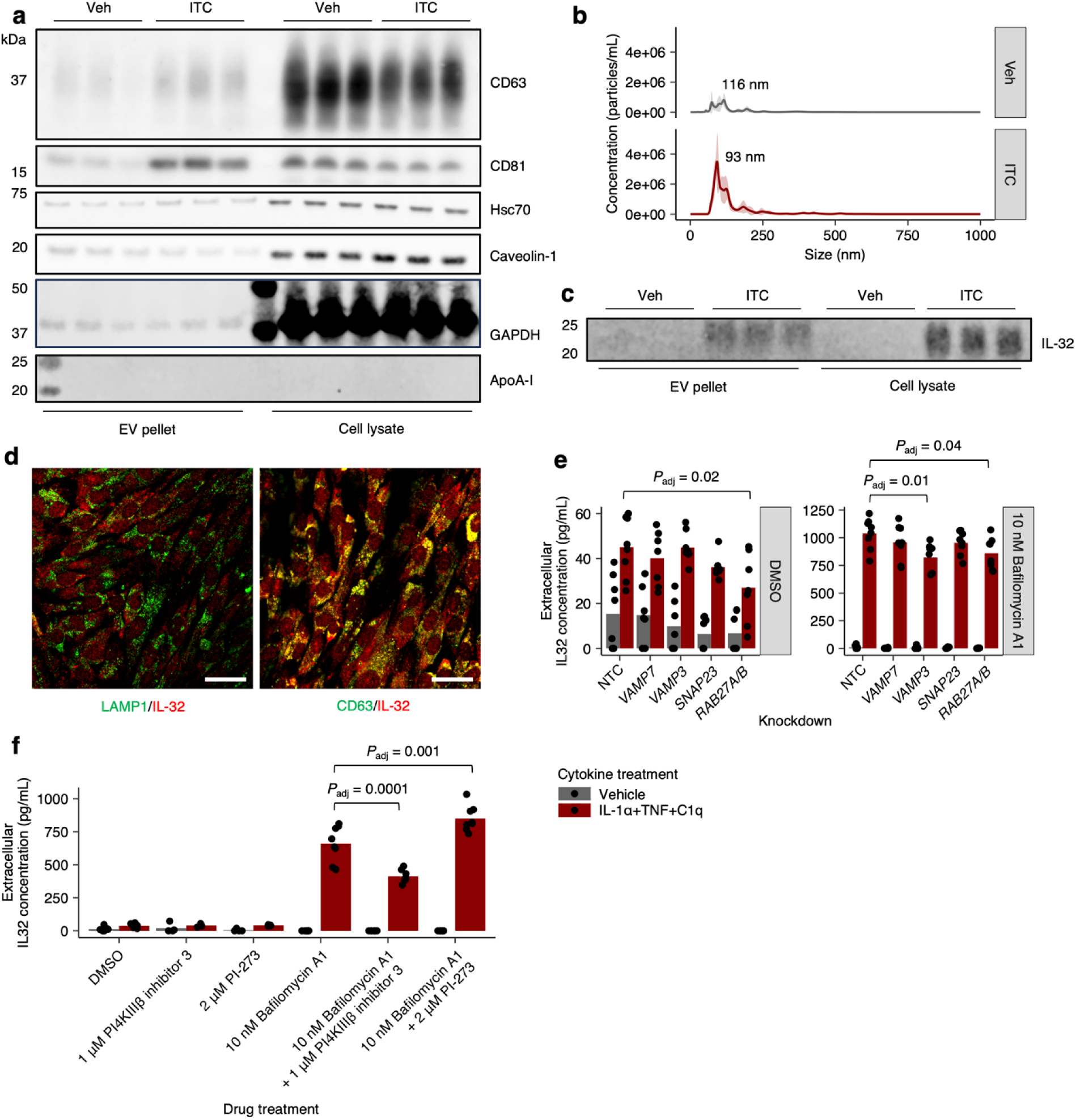
Extracellular IL-32 co-fractionates with extracellular vesicles likely derived from multivesicular body exocytosis. **a**, Immunoblots against consensus extracellular vesicle (EV) markers (CD63, CD81, Hsc70, Caveolin-1) or common contaminants (e.g. ApoA-I) in EVs isolated from iAstrocyte conditioned media or total cell lysate after vehicle vs. ITC treatment. **b**, EV size distribution measured by nanoparticle tracking analysis. **c**, Immunoblot against IL-32 in EVs isolated from iAstrocyte conditioned media or total cell lysate. **d**, Representative images of dual immunostaining against LAMP1 together with IL-32 or CD63 together with IL-32; scale bar = 60 μm. **e**-**f**, Extracellular IL-32 concentration measured by ELISA in conditioned media from ITC- vs. vehicle-treated iAstrocytes transduced with non-targeting (NTC) sgRNAs or sgRNAs targeting genes encoding proteins involved in multivesicular body exocytosis (**e**), or treated with small molecules known to inhibit (PI4KIIIβ inhibitor 3) or not inhibit (PI-273) exosome biogenesis. *P* values were calculated by linear regression with correction for multiple testing by Holm’s method, shown only when significant.

Having validated our EV preparation, we proceeded to blot for IL-32 and found that IL-32 was present in EVs isolated from ITC-treated but not vehicle-treated iAstrocytes (Fig. 5c). To see if extracellular IL-32 was exclusively EV-associated, we analyzed the EV pellet and supernatant by western blotting and ELISA. On western blot, IL-32 was detectable only in the EV pellet, as were EV markers (Supplementary Fig. 7a). However, by ELISA, we saw that extracellular IL-32 partitioned roughly equally between the EV pellet and supernatant (Supplementary Fig. 7b). In comparison, CTSB exclusively partitioned into the supernatant (Supplementary Fig. 7c). The discrepancy between the western blot and ELISA data for IL-32 could potentially be explained by the fact that the western blot and ELISA antibodies are polyclonal antibodies likely raised against different immunogens. Perhaps the ELISA antibodies detect an IL-32 isoform not detected by the western blot antibody.

Since EVs can be derived from direct budding of the plasma membrane or multivesicular body exocytosis (which would then be referred to as exosomes) (52), we performed immunostaining of ITC-treated iAstrocytes and visualized the subcellular localization of IL-32 with confocal microscopy. We found that IL-32 co-localized strongly with CD63 (Fig. 5d, Supplementary Fig. 8), which is also a marker of multivesicular bodies (53), suggesting that extracellular IL-32 is derived from multivesicular body exocytosis. We proceeded to further test this hypothesis by observing the effect of bafilomycin A1 or knockdown of genes encoding proteins involved in multivesicular body exocytosis on extracellular IL-32 levels. In agreement with prior reports which demonstrated increased exosome release with bafilomycin A1 (34,36,54), we found that treatment with bafilomycin A1 dramatically increased extracellular IL-32 levels (Fig. 5e). Furthermore, knockdown of *RAB27A/B* decreased extracellular IL-32 levels (Fig. 5e), consistent with the known role of RAB27A/B in exosome secretion (55). Lastly, corroborating a prior report which identified a role for PI4KIIIβ in exosome biogenesis (54), we found that inhibition of PI4KIIIβ (using PI4KIIIβ inhibitor 3 (56)) but not PI4KIIα (using PI-273 (57)) decreased extracellular IL-32 levels in the presence of bafilomycin A1 (Fig. 5f).

Having explored the secretion mechanism of IL-32, we subsequently characterized its function in inflammatory astrocyte reactivity. Our prior work established two distinct polarizations of inflammatory reactive astrocytes after ITC treatment – a VCAM1^+^ interferon/TNF-responsive polarization associated with CXCL10 secretion vs. a VCAM1^-^/C3^+^ IL-1/IL-6-responsive polarization associated with GM-CSF secretion (9). After verifying that knockdown of *IL32* resulted in robust depletion of IL-32 at the protein level (Supplementary Fig. 9a-b), we found that *IL32* knockdown subtly decreased the proportion of VCAM1^+^/C3^-^ astrocytes and noticeably decreased CXCL10 secretion (Fig. 6a-b), suggesting that IL-32 promotes the interferon/TNF-responsive polarization. Also, IL-32 induction by ITC was greater in VCAM1^+^/C3^+^ interferon-responsive astrocytes compared to IL-1/IL-6-rresponsive VCAM1^-^/C3^+^ astrocytes (Supplementary Fig. 9c-e). Furthermore, we found that IFN-β increased the upregulation of IL-32 by ITC (Supplementary Fig. 9f), consistent with prior work demonstrating a role of IL-32 in antiviral responses (58–60).

**Figure 6.**
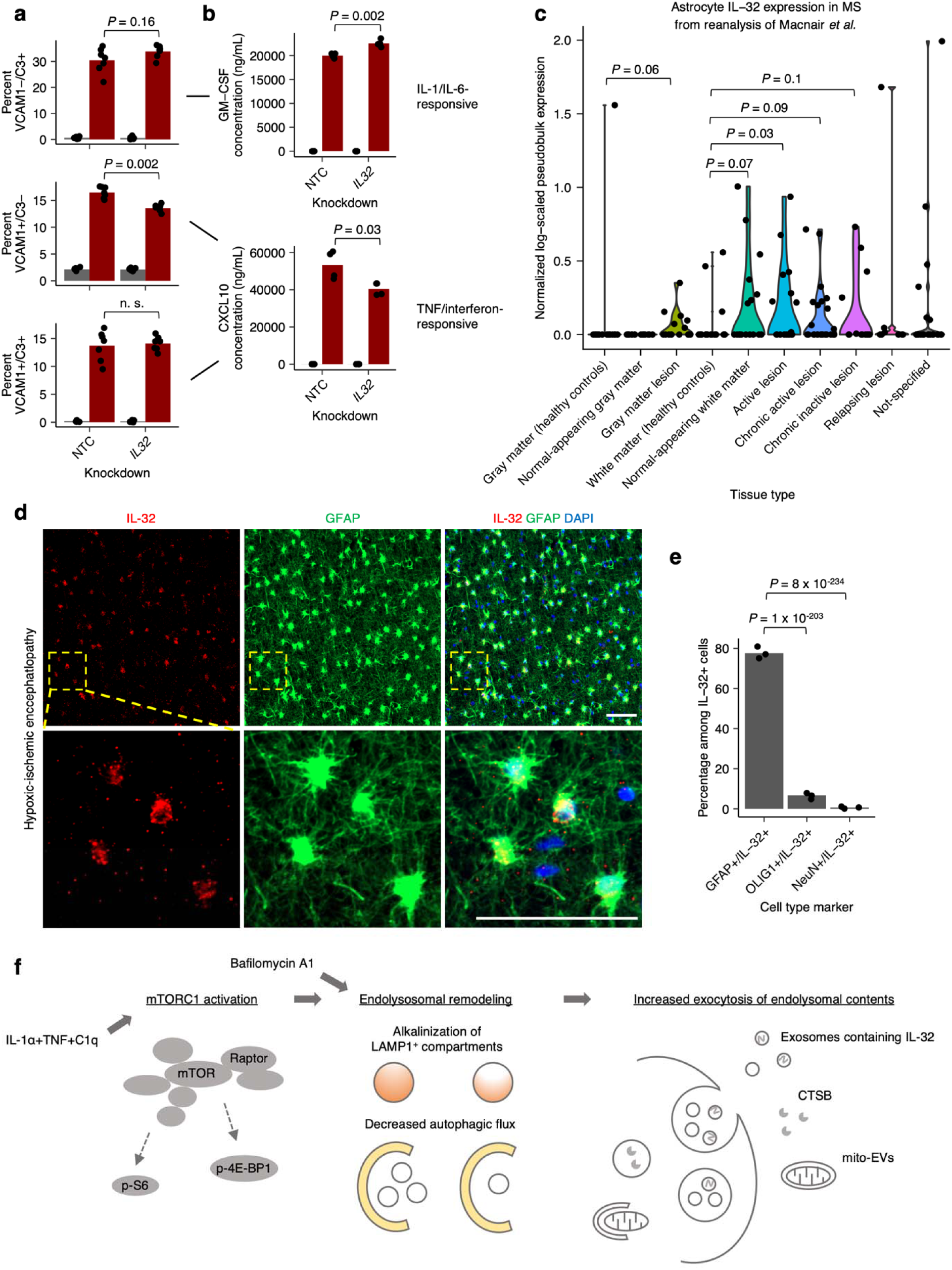
IL-32 regulates the polarization of inflammatory reactive astrocytes and is upregulated in astrocytes in neuroinflammatory conditions. **a**-**b**, Proportion of IL-1/IL-6-responsive (VCAM1-/C3+) or TNF/interferon-responsive (VCAM1+/C3-, VCAM1+/C3+) inflammatory reactive astrocyte polarizations (**a**) or their associated cytokines (**b**) in ITC- vs. vehicle-treated iAstrocytes transduced with non-targeting (NTC) sgRNAs or sgRNAs targeting *IL32*. **c**, Log-scaled *IL32* expression in astrocytes found in normal tissue vs. multiple-sclerosis lesions derived from pseudobulk analysis of snRNA-seq data from Macnair *et al.*; n = 15 for healthy control gray matter, n = 15 for normal-appearing gray matter, n = 15 for gray matter lesion, n = 22 for healthy control white matter, n = 18 for normal-appearing white matter, n = 17 for active lesion, n = 27 for chronic active lesion, n = 13 for chronic inactive lesion, n = 8 for relapsing lesion, n = 23 for not specified. **d**, Representative immunostaining of IL-32 and GFAP in white matter brain tissue from patients with hypoxic-ischemic encephalopathy (HIE); scale bar 50 μm. e, Percent GFAP+, OLIG1+, or NeuN+ cells among IL-32+ cells in HIE brain tissue (n = 3 patients); *P* values calculated via beta regression. **f**, Schematic of ITC-induced, mTOR-dependent endolysosomal remodeling and associated exocytic activity. *P* values were calculated using the Mann-Whitney U test in **a**, **c**, and **e**, and using the two-sided Student’s t test in **b**.

Given our finding that extracellular IL-32 is associated with EVs, we wondered if free extracellular IL-32 could influence inflammatory reactive astrocyte states. We found that treatment with recombinant IL-32β or IL-32γ at 200 ng/mL did not appreciably alter the proportion of VCAM1^+^/C3^-^ astrocytes, regardless of *IL32* knockdown (Supplementary Fig. 9f). As the concentration of IL-32β and IL-32γ we used here is well above the range where immune cells could be robustly activated by recombinant IL-32 (61), our data suggests that the effect of IL-32 on inflammatory reactive astrocytes states may be mediated intracellularly, either through uptake of IL-32-containing EVs or cell-autonomously through IL-32 that has not been secreted.

Next, given that IL-32 levels have been reported to be elevated in the cerebrospinal fluid of patients with multiple sclerosis and neuro-Behcet’s disease (15), we wanted to see if IL-32 was upregulated in astrocytes under neuroinflammatory conditions. Although the fact that IL-32 does not have an ortholog in rodents precluded us from analyzing the vast amount of published transcriptomic data on mouse models of neuroinflammation, we found that *IL32* transcript levels were indeed upregulated in astrocytes in various types of multiple sclerosis lesions in humans (Fig. 6c) by reanalyzing the comprehensive single-nucleus RNA-seq dataset from Macnair *et al.* (62) (121 subjects total) at the pseudobulk level (63) (see Methods). There was also evidence of *IL32* upregulation in oligodendrocytes, microglia, and endothelial cells, but only in white matter lesions (Supplementary Fig. 10). Interestingly, astrocytes were the only cell type with evidence of *IL32* upregulation in gray matter lesions (Fig. 6c). To examine whether IL-32 could be found in other neuroinflammatory conditions, we stained for IL-32 and cell type markers in post-mortem brain tissue from pediatric patients diagnosed with hypoxic-ischemic encephalopathy, a condition involving significant neuroinflammation (63) where we had previously identified upregulation of inflammatory reactive astrocyte markers (9). We found that IL-32 preferentially colocalized with GFAP^+^ astrocytes and to a much lesser degree with OLIG1+ oligodendrocytes (Fig. 6d-e). IL-32 was not detected in NeuN+ neurons (Fig. 6e), consistent with our observations from the multiple sclerosis snRNA-seq data.

Finally, to ensure the overall robustness of our results, we validated key findings in iAstrocytes derived from an independent hiPSC line of different sex (162D). We confirmed that after ITC treatment, 162D iAstrocytes upregulated cell-surface LAMP1 (Supplementary Fig. 11a), accumulated LC3 and p62 puncta (Supplementary Fig. 11b-c), upregulated mTORC1 activity as measured by phospho-S6 staining (Supplementary Fig. 11d), upregulated IL-32 (Supplementary Fig. 11e), and secreted a greater number of mito-EVs (Supplementary Fig. 11h). Furthermore, we also directly characterized lysosome function by assaying intracellular CTSB activity in both 162D iAstrocytes and WTC11 iAstrocytes; ITC treatment decreased intracellular CTSB activity in both 162D and WTC11 iAstrocytes (Supplementary Fig. 11g).

## DISCUSSION

Our results establish mTOR activation as a key feature of inflammatory astrocyte reactivity induced by IL-1α+TNF+C1q (ITC), driving endolysosomal remodeling manifesting as alkalinization of LAMP1^+^ compartments and reduced autophagic flux (Fig. 6e). While we have focused on mTOR-dependent endolysosomal remodeling driven by acute ITC treatment, other cellular processes may also contribute to endolysosomal remodeling. For example, cellular senescence is associated with a profound remodeling of lysosome function and content (64), which in fact may partially occur through mTORC1 hyperactivation (65). Although acute treatment with inflammatory cytokines is unlikely to induce cellular senescence, we nevertheless found overlap of senescence-associated genesets with genes upregulated by ITC (Supplementary Fig. 2b). We speculate that perhaps chronic inflammatory activation of astrocytes could lead to cellular senescence which would further contribute to endolysosomal remodeling.

Connecting our results here with the broader literature on phenotypes associated with inflammatory astrocyte reactivity, we suspect that mTOR-induced endolysosomal remodeling may account for the loss of phagocytic activity observed in inflammatory reactive astrocytes (14). Although we did not measure phagocytic activity here, we observed in our previously published CRISPRi screens on inflammatory reactivity that *MTOR* knockdown rescued the phagocytic deficit induced by ITC (9).

Here, we found that mTOR-dependent endolysosomal remodeling also resulted in the increased exocytosis of certain endolysosomal cargos (Fig. 6e), with increased cell-surface LAMP1 likely acting as a non-specific marker of endolysosomal exocytic activity. Interestingly, LAMP1^+^ astrocytes have been shown to modify disease progression in EAE (64). Furthermore, all three of the endolysosomal cargos we have characterized – CTSB, IL-32, and mito-EVs – have been reported to be involved in neuroinflammatory conditions. CTSB levels are elevated in the CSF as well as brain parenchyma of patients with Alzheimer’s disease (65–68), and knockout of *Ctsb* has been shown to ameliorate the neuropathology and behavioral deficits in mouse models of Alzheimer’s disease (69–71). Mito-EVs (72) have been shown to mediate the transfer of mitochondria from astrocytes to neurons after experimentally induced stroke in mice (48). Lastly, IL-32 levels are elevated in the cerebrospinal fluid of patients with multiple sclerosis or neuro-Behcet’s disease (15), and a polymorphism in the IL-32 promoter has been associated with increased risk of multiple sclerosis in two independent studies (73,74).

Given that IL-32 does not have an ortholog in rodents and that its secretion mechanism is still incompletely characterized, we focused on elucidating IL-32 secretion, taking advantage of our in vitro hiPSC-derived astrocyte platform. We found that after ITC treatment, intracellular IL-32 colocalized with multivesicular bodies, and that extracellular IL-32 co-fractionated with EVs. Whether extracellular IL-32 is exclusively associated with EVs remains to be seen, as we obtained conflicting data by western blotting vs. ELISA. Given that the western blot and ELISA antibodies are polyclonal antibodies likely raised against different immunogens, it is possible that the ELISA antibodies recognize a non-EV-associated IL-32 isoform not detected by the western blot antibody. Alternatively, if extracellular IL-32 consists predominantly of one isoform, it may be cleaved in a way such that the western blot antibody recognizes only the EV-associated fragment whereas the ELISA antibodies recognize both the EV-associated and free fragments. Further biochemical experiments will be necessary to elucidate whether IL-32 isoforms may be differentially associated with EVs or if extracellular cleavage of IL-32 occurs.

With respect to the secretion mechanism of extracellular IL-32, we found that knockdown of genes encoding proteins involved in multivesicular body exocytosis such as RAB27A/B decreased extracellular levels of IL-32, as did pharmacological inhibition of exosome biogenesis. Overall, our results corroborate previous reports demonstrating that a portion of extracellular IL-32 is vesicle-associated (50,51), and we establish exosomes as the likely candidate. With respect to extracellular IL-32 not contained within EVs, it is possible that IL-32 may be loaded directly into multivesicular body lumens and subsequently exocytosed, or that intracellular IL-32 may be released directly through plasma membrane leakage in dying or dead cells (75).

As for the role of IL-32 in neuroinflammation, we found that knockdown of *IL32* affected the polarization of inflammatory reactive states induced by ITC (9), decreasing the abundance of the interferon/TNF-responsive state and its associated cytokine CXCL10. A limitation of knocking down *IL32* with DNA-targeting CRISPRi (76) is that we could not distinguish the contribution of the many splice isoforms of IL-32, which have been reported to have distinct activities (77); future work could elucidate this using RNA interference or RNA-targeting CRISPR-based systems (78).

Regarding how IL-32 acts upon cells, it is an open question whether extracellular IL-32 signals through cell-surface receptors or exerts its effects intracellularly, for example, after EV-mediated uptake or cell-autonomously when it is not secreted. Depending on the cell type and biological context, there is evidence for both extracellular and intracellular activity (16). In our hands, free extracellular recombinant IL-32 at concentrations capable of activating immune cells (61) did not appreciably influence inflammatory reactive astrocyte states, suggesting that our IL-32-associated phenotypes may be mediated intracellularly.

In addition to its effects on the polarization of inflammatory reactive astrocyte states, we also found that IL-32 was induced by IFN-β (an old disease-modifying treatment for multiple sclerosis), and that *IL32* transcript levels were upregulated in astrocytes in various multiple sclerosis lesions. Considering the human genetics data demonstrating the importance of IL-32 to the pathogenesis of multiple sclerosis, studying how astrocyte IL-32 contributes to multiple sclerosis would be a worthwhile future research direction. Lastly, we found preferential colocalization of IL-32 with astrocytes in a different neuroinflammatory condition – hypoxic-ischemic encephalopathy (HIE), suggesting that IL-32 may play a role in HIE as well.

In conclusion, we believe that our results highlight mTOR-dependent endolysosomal remodeling as an important and previously underappreciated aspect of inflammatory astrocyte reactivity which can be targeted therapeutically. We also clarified the secretion mechanism and functional role of an important disease-associated cytokine, IL-32, in astrocytes, a cell type in which IL-32 has rarely been studied (79). Since the receptor for IL-32 is still unknown (16), we believe that our results establish a strong foundation for future studies focused on the how IL-32 mediates its biological effects and contributes to neuroinflammation.

## DECLARATIONS

### Ethics approval and consent to participate

N/A

### Consent for publication

N/A

### Availability of data and materials

The source data and code used to analyze the data presented in this study will be shared upon request. The source data for all western blots shown in this study can be found in Supplementary File 1.

### Author contributions

K.L., B.R., and M.K. conceptualized and led the overall project, and wrote the manuscript with input from all co-authors. K.L. and B.R. performed the experiments and data analysis unless stated otherwise. In consultation with J.E., F.M aided with sample preparation for whole-cell and lysosome-specific proteomics experiments, and performed mass spectrometry and preliminary computational analysis. I.V.L.R. performed sample prep for the cell-surface LAMP1 CRISPRi screens. K.A.H. guided TIRF experiments. M.L. and S.B aided with design and sample preparation for whole-cell and lysosome-specific proteomics experiments, respectively. M.Y.C. and A.W.K provided FIRE-pHLy reagents and guidance on lysosome pH experiments. S.F conducted LAMP2 immunofluorescence experiments and data analysis. All authors reviewed the manuscript.

## Acknowledgements

We thank Greg Mohl, Avi Samelson, Olivia Teter, Biswa Ramani, Brandon Desousa, Emmy Li, Vukasin Jovanovic, Zuzana Krejciova, Nawei Sun, and Carlo Condello for contributions to preliminary studies or helpful discussions. We thank members of the Kampmann lab for discussions and feedback on the manuscript. We thank the staff at the UCSF Center for Advanced Light Microscopy and the UCSF Parnassus Flow Cytometry CoLab (RRID:SCR_018206) for their assistance. A portion of the imaging data presented in this manuscript were acquired on microscopes at the UCSF Center for Advanced Light Microscopy and the Innovation Core at the Weill Institute, a part of the UCSF Center for Advanced Light Microscopy.

## FIGURES

**Supplementary figure 1.**
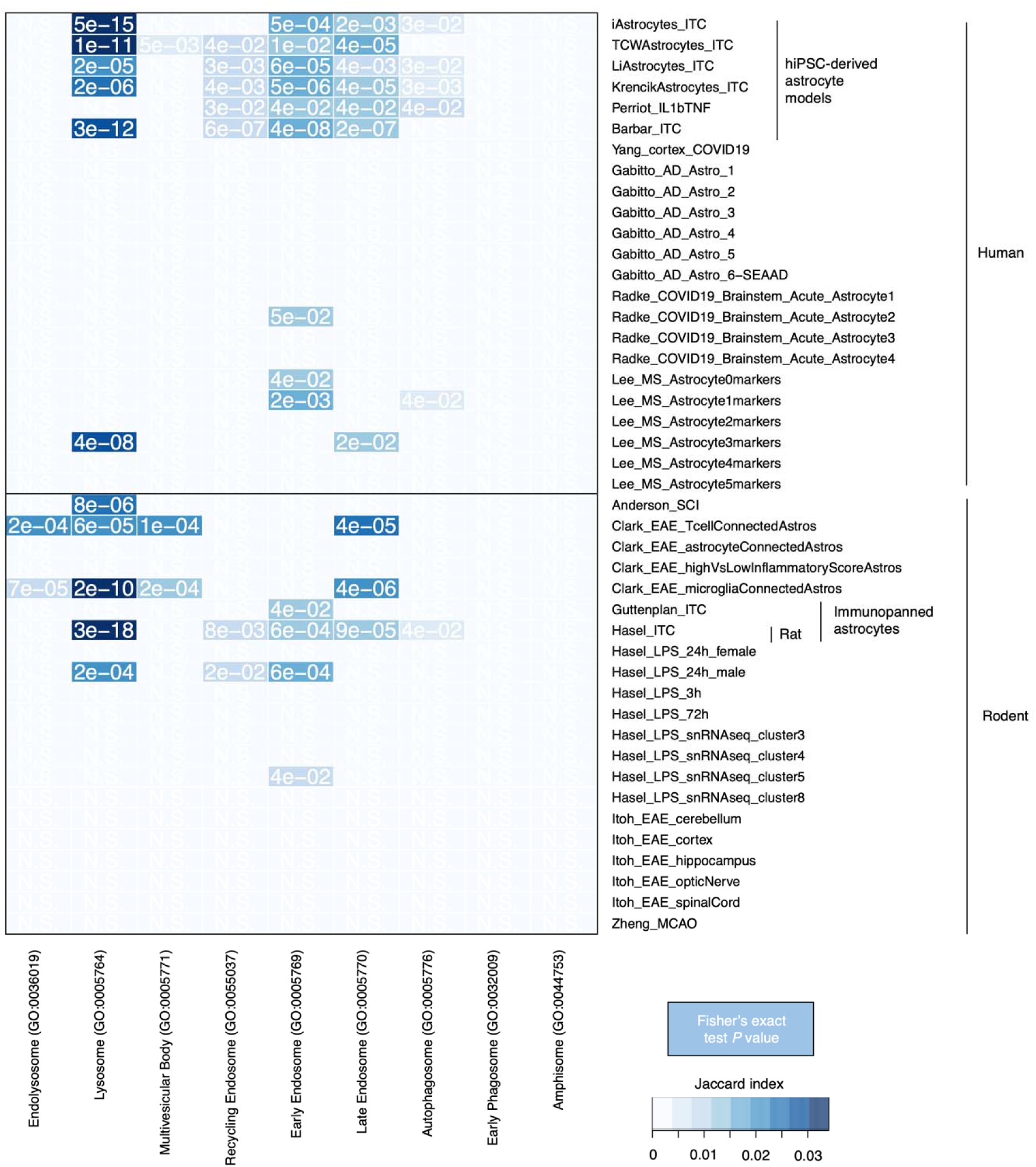
Overlap analysis of downregulated genes with endolysosomal-related terms in inflammatory reactive astrocytes across different contexts and species. GO Cellular Component terms corresponding to different components of the endolysosomal system were analyzed for overlap with downregulated genes in human or rodent astrocytes (all from mouse except where noted) found in various neuroinflammatory models and conditions. See Supplementary Table 3 for metadata corresponding to the studies analyzed. *P* values were adjusted for multiple comparisons with the Benjamini-Hochberg procedure.

**Supplementary figure 2.**
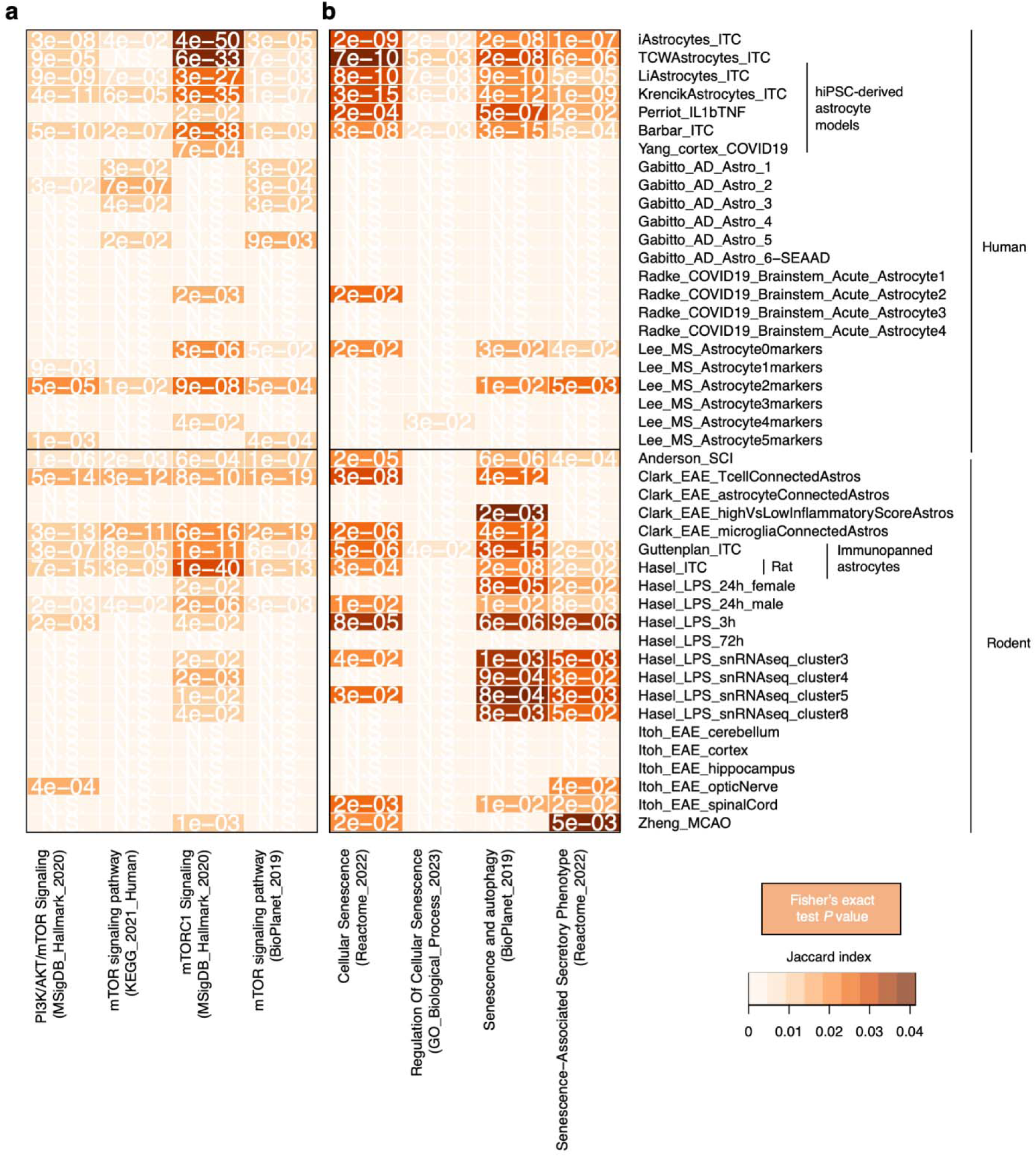
Overlap analysis of upregulated in inflammatory reactive astrocytes across different contexts and species. mTOR-related terms (**a**) or cellular senescence-related terms (**b**) were analyzed for overlap with upregulated genes in human or rodent astrocytes (all from mouse except where noted) found in various neuroinflammatory models and conditions. See Supplementary Table 3 for metadata corresponding to the studies analyzed. *P* values were adjusted for multiple comparisons with the Benjamini-Hochberg procedure.

**Supplementary figure 3.**
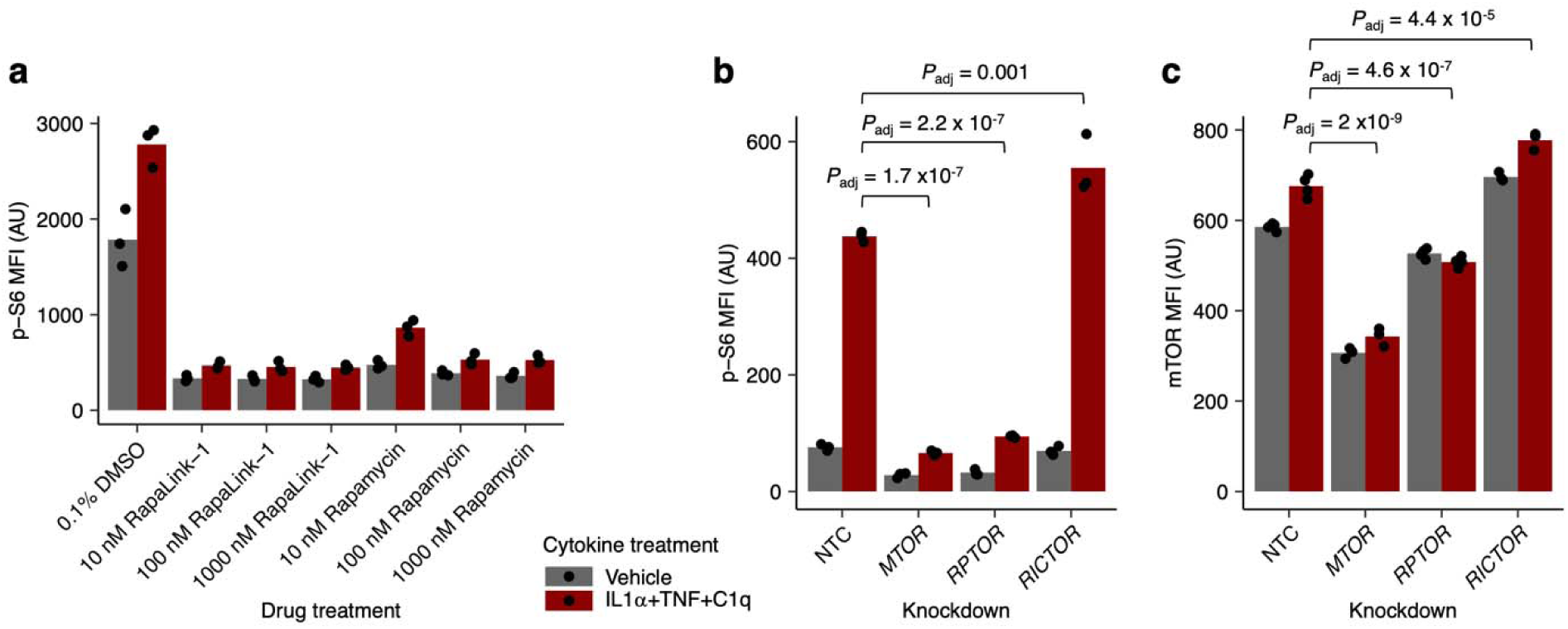
Validation of mTOR inhibition and knockdown. **a**, Median fluorescence intensity (MFI) of phospho-S6 measured by flow cytometry in ITC- vs. vehicle-treated iAstrocytes co-treated with rapamycin or Rapalink-1. **b**-**c**, Phospho-S6 (**b**) or mTOR (**c**) MFI measured by flow cytometry in vehicle- or ITC-treated iAstrocytes transduced with a non-targeting sgRNA (NTC) or sgRNAs targeting genes encoding mTORC1/2 subunits; n = 3 wells for all conditions; *P* values calculated by linear regression with Holm’s correction for multiple testing, shown for selected comparisons).

**Supplementary figure 4.**
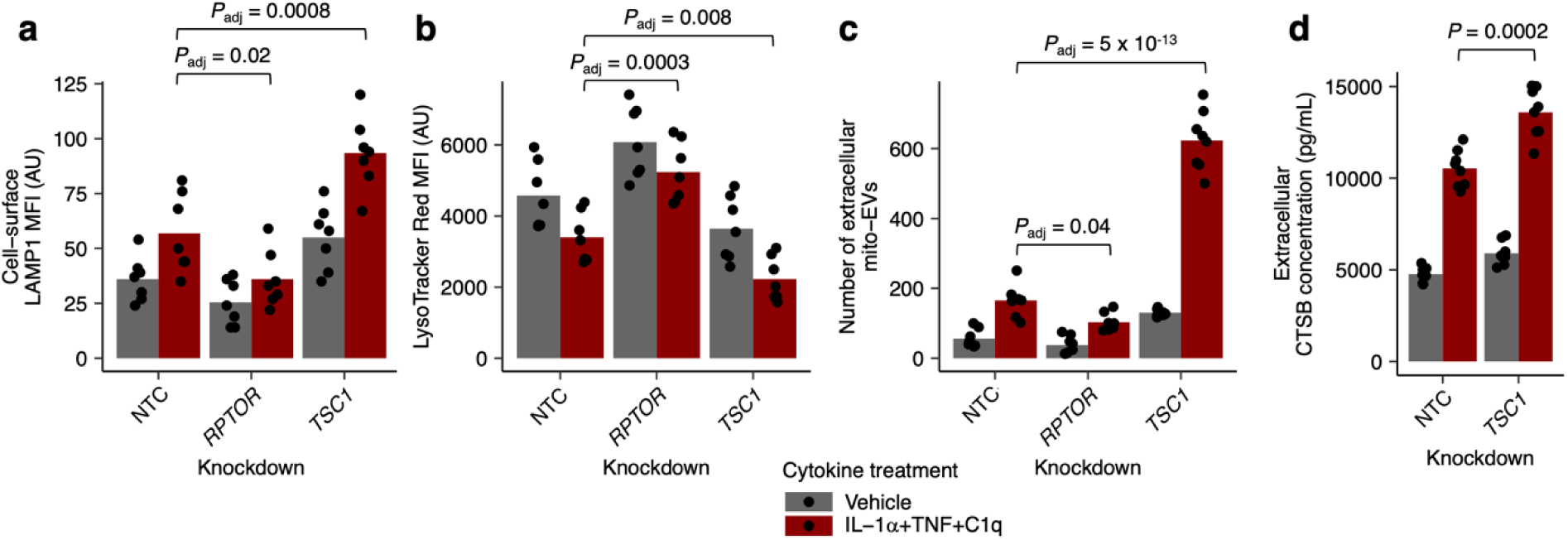
Modulation of mTORC1 activity affects endolysosomal exocytosis bidirectionally. **a**-**d**, Median fluorescence intensity (MFI) of cell-surface LAMP1 (**a**) or LysoTracker (**b**) staining measured by flow cytometry, abundance of mito-EVs in conditioned media (**c**), or extracellular CTSB concentration measured by electrochemiluminescence-based immunoassay (**d**) in ITC- vs. vehicle-treated iAstrocytes transduced with non-targeting (NTC) sgRNAs or sgRNAs targeting *RPTOR* or *TSC1* (n = 7 wells per condition for LAMP1 and LysoTracker data, n = 8 wells per condition for mito-EV and CTSB data; *P* values calculated by linear regression with Holm’s correction for multiple testing, shown only if significant).

**Supplementary figure 5.**
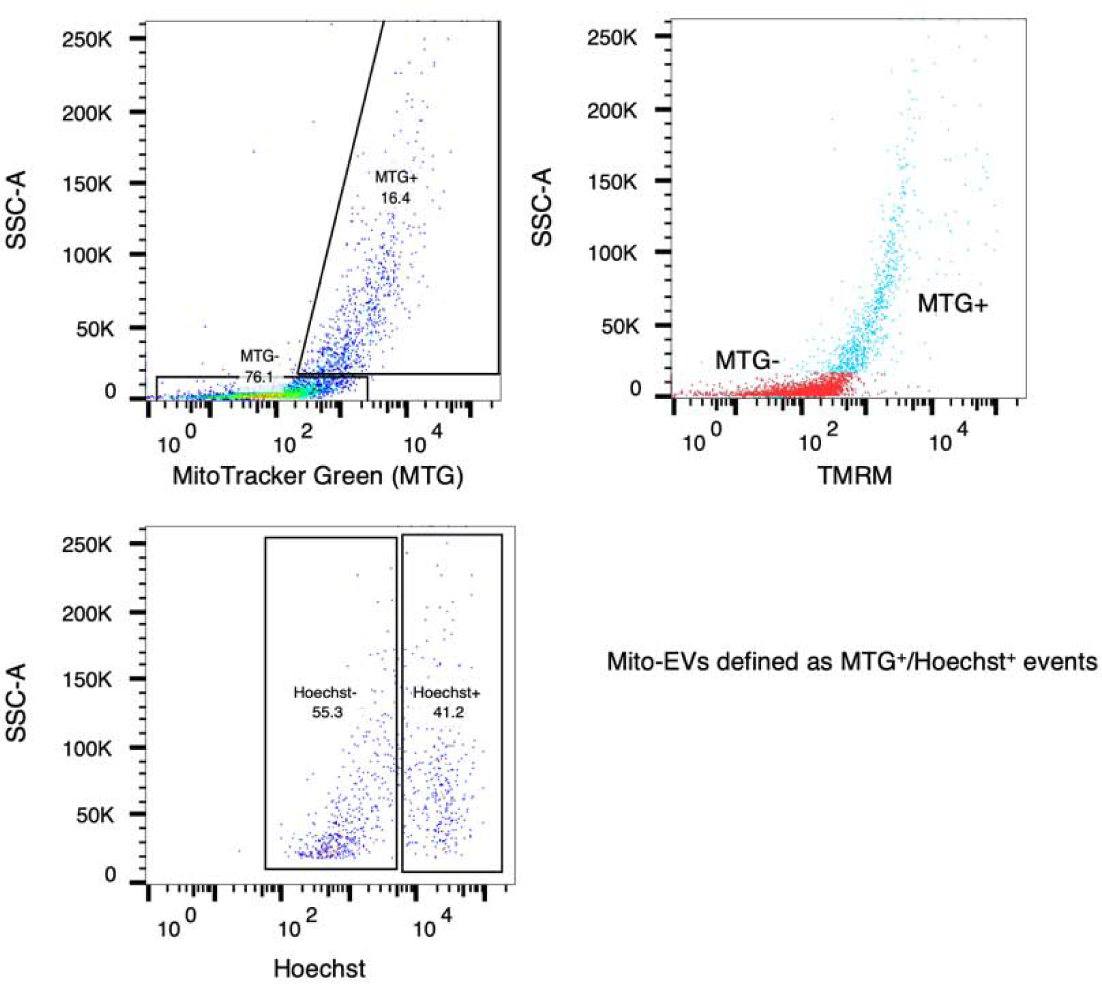
Gating strategy for quantifying mito-EVs from iAstrocyte conditioned media. Events were first gated on MitoTracker Green fluorescence, with MTG^+^ events subsequently gated by Hoechst staining. Mito-EVs were defined as MTG^+^/Hoechst^+^ events.

**Supplementary figure 6.**
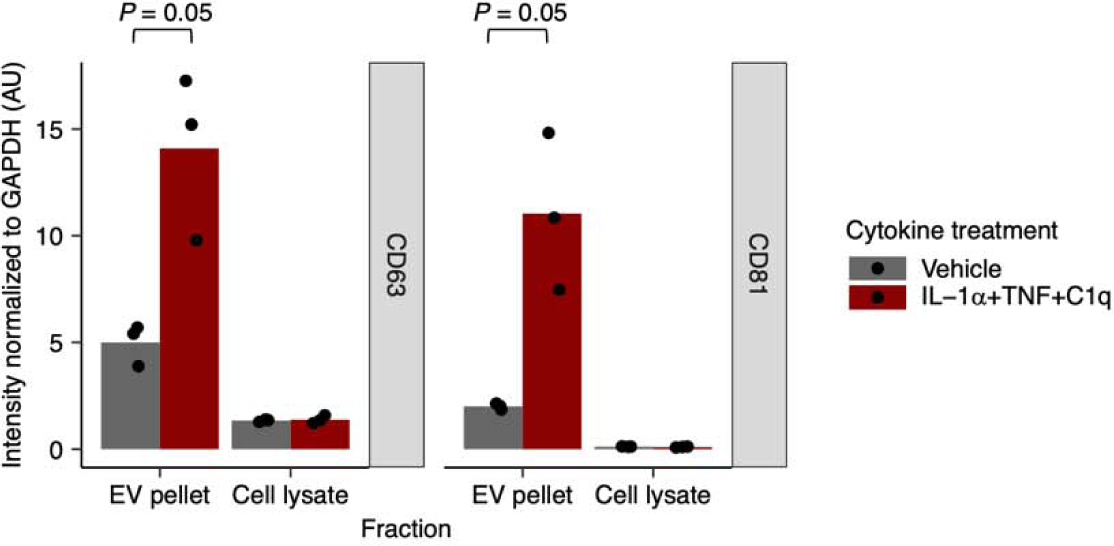
ITC-treated astrocytes release more extracellular vesicles. Quantification of CD63 or CD81 immunoblot bands from total cell lysate or extracellular vesicles from ITC- vs. vehicle-treated iAstrocytes shown in Fig. 5a (n = 3 wells per condition; *P* values by two-sided Student’s t test).

**Supplementary figure 7.**
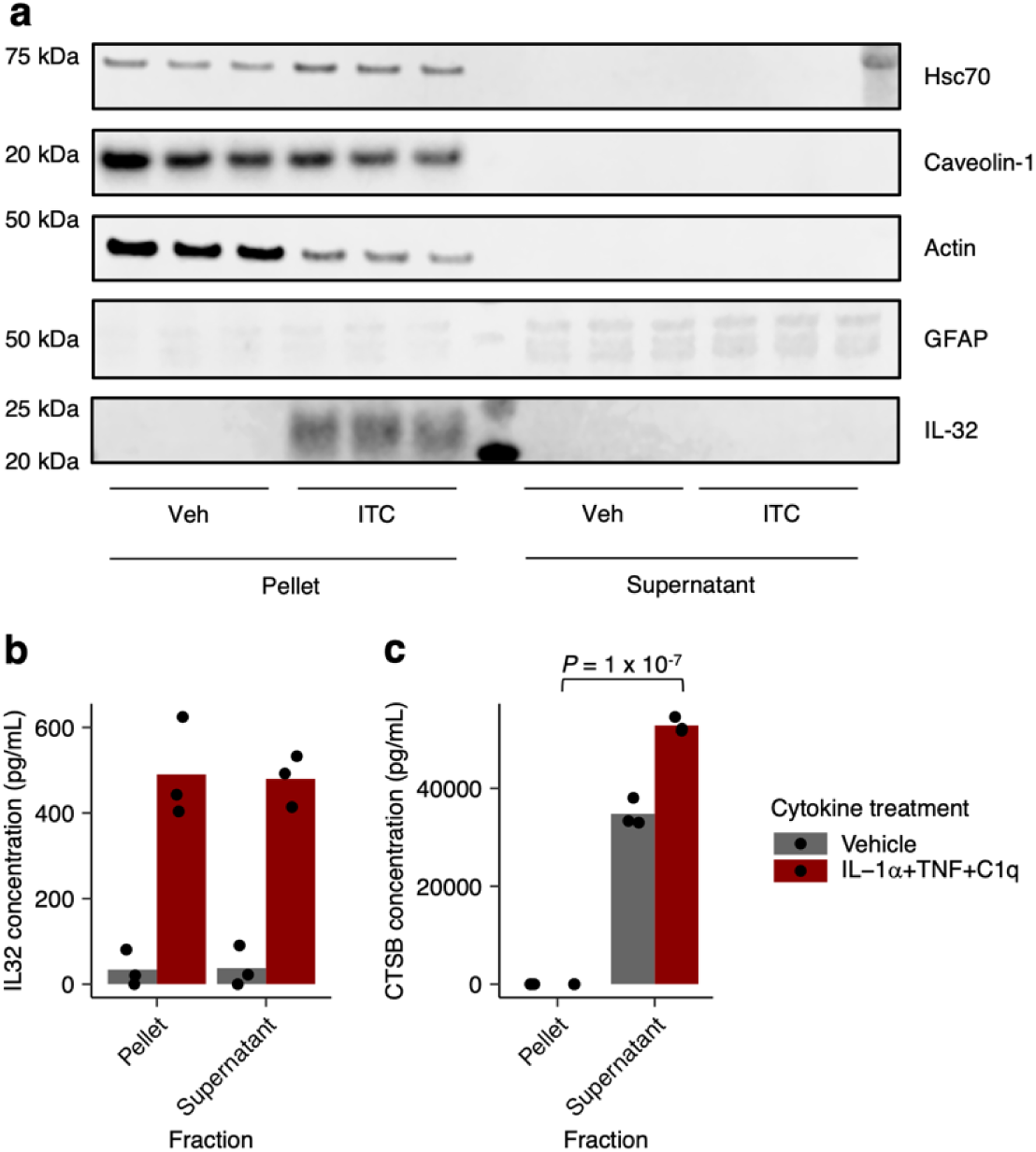
Partition analysis of IL-32 or CTSB. Concentration of IL-32 measured by ELISA (**a**) or CTSB measured by electrochemiluminescence-based immunoassay (**b**) in resuspended extracellular vesicle pellets (lysed with detergent) vs the supernatant (see Methods); n = 3 per condition, *P* values calculated only for ITC-treated conditions by two-sided Student’s t test, shown only when significant.

**Supplementary figure 8.**
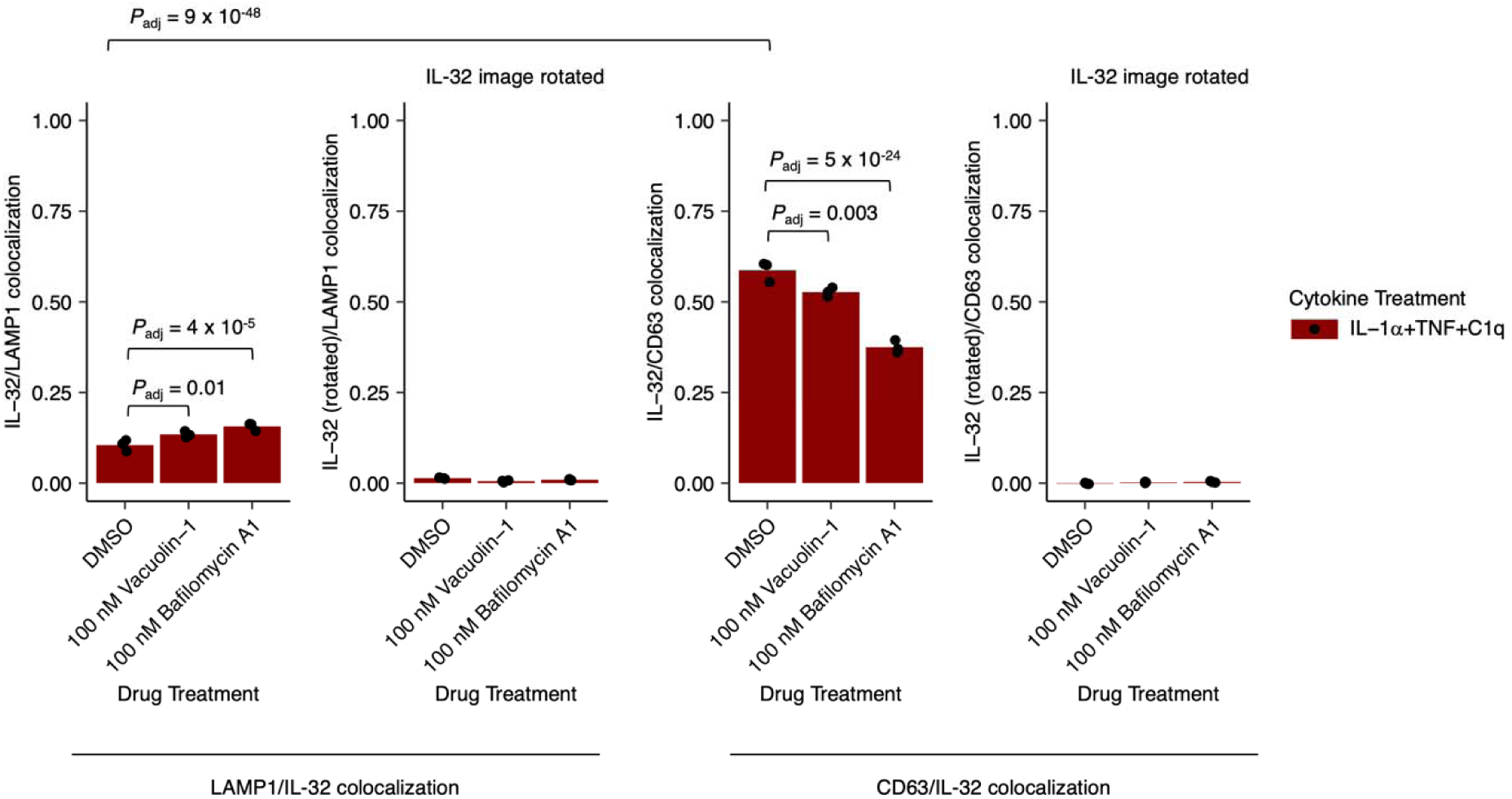
Colocalization analysis of IL-32 with LAMP1 or CD63. Pearson’s colocalization coefficient between IL-32 and LAMP1 (left) or between IL-32 and CD63 (right) in ITC-treated iAstrocytes co-treated with vacuolin-1 or bafilomycin A1, corresponding to images from the immunostaining experiment shown in Fig. 5d (n = 3 wells per condition; *P* values calculated by beta regression with correction for multiple testing by Holm’s method, shown only if significant).

**Supplementary Figure 9.**
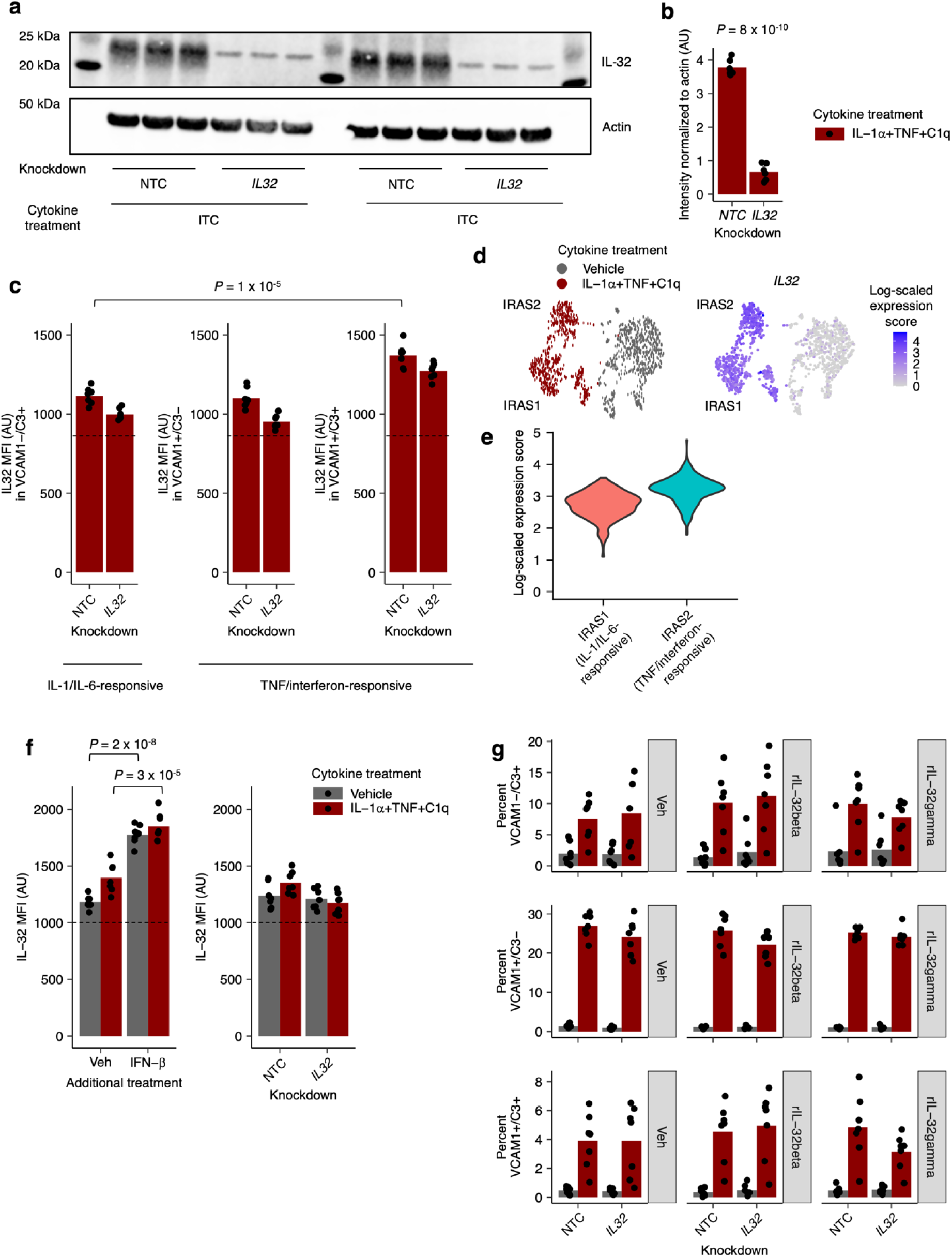
IL-32 levels are higher in interferon/TNF-responsive inflammatory reactive astrocytes and after treatment with IFN-β. **a**, Immunoblot against IL-32 in ITC-treated iAstrocytes transduced with non-targeting (NTC) sgRNAs or sgRNAs targeting *IL32.* b, Quantification of IL-32 levels from immunoblot shown in **a**; n = 6 wells per condition. **c**, Median fluorescence intensity (MFI) of IL-32 in IL-1/IL-6-responsive (VCAM1-/C3+) or TNF/interferon-responsive (VCAM1+/C3-, VCAM1+/C3+) inflammatory reactive astrocytes as defined in Leng *et al.* <u>(Leng et al. 2022)</u>; n = 7 wells per condition. **d**-**e**, *IL32* transcript levels from single-cell RNA-seq data on ITC- vs. vehicle-treated iAstrocytes from Leng *et al.* <u>(Leng et al. 2022)</u> visualized on uniform manifold approximation embedding (**d**) or as violin plots (**e**). **f**, IL-32 MFI in ITC- vs. vehicle-treated iAstrocytes treated concurrently with vehicle or IFN-β transduced with non-targeting (NTC) sgRNAs or sgRNAs targeting *IL32*; n = 7 wells per condition. g, Proportion of VCAM1-/C3+, VCAM1+/C3-, or VCAM1+/C3+ iAstrocytes after vehicle vs. ITC treatment with or without IL-32 knockdown or treatment with free recombinant IL-32β or IL-32γ (200 ng/mL); n = 7 wells per condition. Dotted lines in **c** and **f** reflect estimated level of non-specific staining based on degree of IL-32 knockdown from b. *P* values where shown were calculated by two-sided Student’s t test.

**Supplementary Figure 10.**
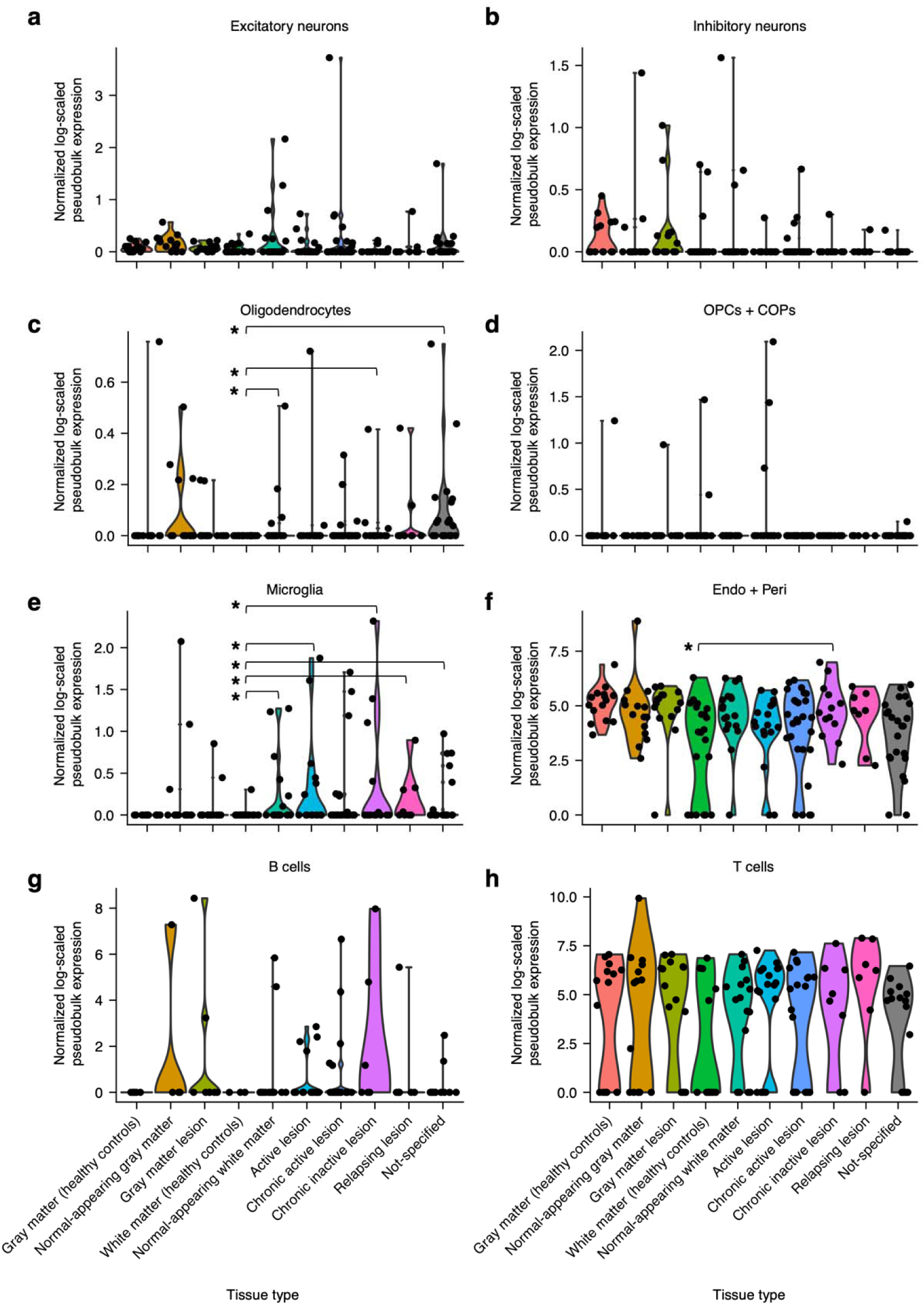
IL32 expression in non-astrocyte cell types in snRNA-seq data from Macnair *et al*. **a**-**f**, Log-scaled normalized pseudobulk counts of *IL32* in excitatory neurons (**a**), inhibitory neurons (**b**), oligodendrocytes (**c**), oligodendrocyte precursor cells (OPCs) and differentiation-committed oligodendrocyte precursors (COPs) (**d**), microglia (**e**), or endothelial cells s(endo) and pericytes (peri) (**f**). *P* values were calculated with the Mann-Whitney U test and were not adjusted for multiple comparisons; * *P* < 0.05. n = 15 for healthy control gray matter, n = 15 for normal-appearing gray matter, n = 15 for gray matter lesion, n = 22 for healthy control white matter, n = 18 for normal-appearing white matter, n = 17 for active lesion, n = 27 for chronic active lesion, n = 13 for chronic inactive lesion, n = 8 for relapsing lesion, n = 23 for not specified.

**Supplementary Figure 11.**
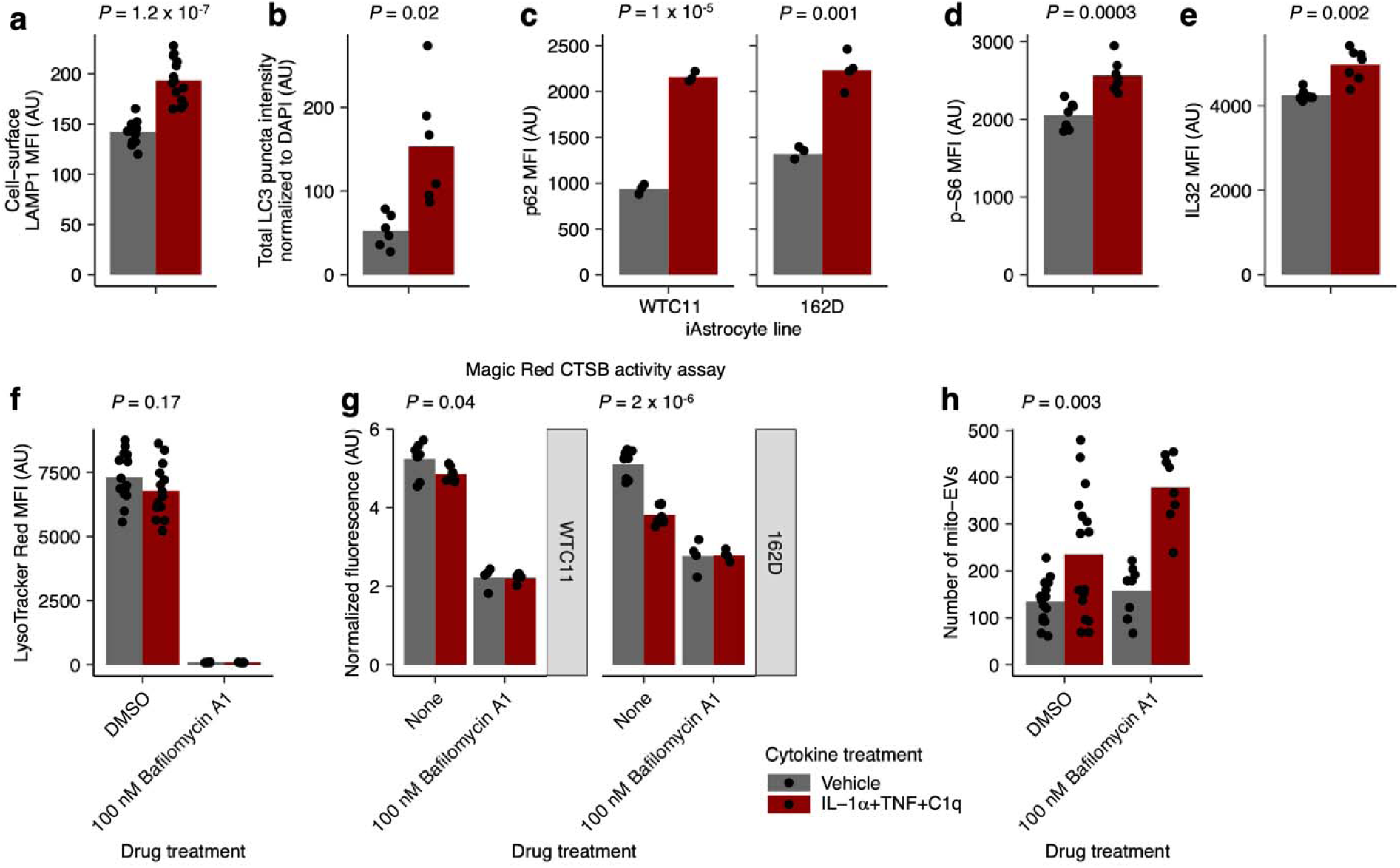
Validation experiments using iAstrocytes derived from the 162D hiPSC line. **a**, Median fluorescence intensity (MFI) of cell-surface LAMP1 measured by flow cytometry in iAstrocytes treated with vehicle vs. ITC; n = 14 wells per condition. **b**, Total intensity of LC3+ puncta measured by immunofluorescence imaging in iAstrocytes treated with vehicle vs. ITC; n = 6 wells per condition. **c**-**e**, MFI of p62 (**c**), phospho-S6 (**d**), or IL-32 (**e**) measured by flow cytometry in iAstrocytes treated with vehicle or ITC; n = 4 wells per condition for p62, n = 7 wells per condition for phospho-S6 and IL-32. **f**, MFI of LysoTracker staining in iAstrocytes treated with vehicle vs ITC with or without bafilomycin A1 co-treatment; n = 14 wells per condition for DMSO, n = 7 wells per condition for bafilomycin A1. **g**, Intracellular CTSB activity assayed using the Magic Red CSTB substrate, measured by fluorescence plate reader, in iAstrocytes treated with vehicle vs. ITC with or without bafilomycin A1 pre-treatment (3 hours) prior to the CTSB activity assay; n = 8 wells per condition in the absence of bafilomycin A1 pre-treatment, n = 4 wells per condition in the presence of bafilomycin A1 pre-treatment. **h**, Number of mito-EVs measured by flow cytometry in conditioned media from iAstrocytes treated with vehicle vs. ITC with or without bafilomycin A1 co-treatment; n = 16 wells per condition for DMSO, 8 wells per condition for bafilomycin A1. *P* values were calculated with the two-sided Student’s t test.

## METHODS

### hiPSC culture

Human iPSCs (male WTC11 background (80), female 162D background (81)) were cultured in Essential 8 (E8) Medium (ThermoFisher Scientific cat. no. A1517001) on BioLite Cell Culture Treated Dishes (ThermoFisher Scientific) coated with Growth Factor Reduced, Phenol Red-Free, LDEV-Free Matrigel Basement Membrane Matrix (Corning cat. no. 356231) diluted 1:100 in DMEM/F12 (ThermoFisher Scientific cat. no. 11330032). Essential 8 Medium was replaced daily. When hiPSC colonies demonstrated mature morphology, the hiPSCs were either clump passaged with EDTA for routine maintenance or dissociated to a near single-cell suspension with Accutase Cell Dissociation Reagent (ThermoFisher Scientific cat. no. A11105-01) for applications requiring cell counting. For clump passaging with EDTA, hiPSCs were washed with Dulbecco’s phosphate buffered saline (DPBS; Milipore Sigma cat. no. D8537) and then incubated with Versene (ThermoFisher Scientific cat. no. 15040066) for 5-7 min at room temperature; the Versene solution was then aspirated and replaced with E8 + 10 nM Y-27632 dihydrochloride ROCK inhibitor (Tocris cat. no. 125410); hiPSC colonies were then gently detached mechanically using a cell scraper, resuspended gently, and passaged at 1:10-1:30 dilution in E8 + Y-27632, with Y-27632 removed the next day. For near single-cell dissociation, hiPSCs were washed with DPBS, incubated with Accutase for 5-10 min at 37 °C, and then gently triturated with a P1000 pipette tip; the cell suspension was then diluted with PBS, collected into conical tubes, and spun down at 300 g for 3 min; hiPSCs were then resuspended in E8 + Y-27632, counted, and plated onto Matrigel-coated plates at the desired density in E8 + Y-27632; Y-27632 would be maintained until the hiPSC colonies reached the appropriate size (> cells). Studies with hiPSCs at UCSF were approved by the The Human Gamete, Embryo and Stem Cell Research (GESCR) Committee. Informed consent was obtained from the human subject when the WTC11 line was originally derived.

### Generation of iAstrocytes

iAstrocytes were generated as detailed in Leng *et al*. (9). Briefly, we generated hiPSCs (male WTC11 background and female 162D background) used for iAstrocyte differentiation by inserting a CRISPRi machinery cassette (pC13N-dCas9-BFP-KRAB, Addgene plasmid no. 127968) into the CLYBL safe-harbor locus and a doxycycline-inducible *NFIA* and *SOX9* cDNA cassette into the AAVS1 safe-harbor locus using TALEN-based editing. After recovering a monoclonal CRISPRi-*NFIA*-*SOX9* line, we differentiated hiPSCs to the neuroectoderm lineage via an embryoid body (EB)-based neural induction protocol. After dissociating plating EB-derived cells, we selected and expanded those forming neural rosettes. We used fluorescence activated cell sorting (FACS) to enrich for pure CD133^+^/CD271^-^ neural progenitor cell (NPC) populations for expansion. For iAstrocyte differentiation, these NPCs were re-plated on 1:200 Matrigel-coated dishes and cultured with ScienCell Astrocyte Media (ScienCell Research Laboratories cat. no. 1801) + 2 μg/mL doxycycline (Millipore Sigma cat. no. D9891) to initiate iAstrocyte differentiation. A full media change was completed every other day (maintaining doxycycline at 2 μg/mL) until NPCs reached confluency, at which time they were dissociated with Accutase and split 1:10 onto new Matrigel coated dishes (or cryopreserved) for continued differentiation and expansion. Expansion was continued until day 20 (d20) of differentiation, yielding iAstrocytes. See Leng *et al*. for full details of iAstrocyte hiPSC generation and differentiation protocols.

### Induction of inflammatory reactivity in iAstrocytes

For all iAstrocyte experiments in this study, d20 iAstrocytes were plated at 20,000 cells/cm^2^ on 1:200 Matrigel-coated BioLite Cell Culture Treated Dishes (ThermoFisher Scientific) in ScienCell Astrocyte Media (without doxycycline) on d0. Full media changes were completed on d1, d3, and d5. On d5, media was supplemented with IL-1α (3 ng/mL; Peprotech cat. no. AF-200-01A), TNF (30 ng/mL; Peprotech cat. no. AF-300-01A), and C1q (400 ng/mL; Complement Technology cat. no. A099) to induce inflammatory reactivity according to Liddelow *et al*. (14). All experiments were conducted 24 hours after cytokine treatment (day 6 after plating). For experiments involving addition of additional IFN-β, 5 ng/mL of IFN-β was added concurrently with ITC. For experiments involving addition of recombinant IL-32, 200 ng/mL of recombinant IL-32β (R&D Systems cat. no. 6769-IL) or IL-32γ (R&D Systems cat. no. 4690-IL/CF) was added concurrently with ITC.

### CRISPRi-mediated gene knockdown using lentiviral transduction of individual sgRNAs

For knocking down target genes with a single sgRNA, we cloned CRISPRi sgRNAs into pMK1334 (82) (Addgene cat. no. 127965), as previously described (83). For knocking down target genes using two sgRNAs to achieve greater degree of knockdown, we cloned sgRNAs into pMK1334 with an additional hU6 insert as described in Replogle *et al.* (84). We generated lentivirus of the resulting constructs by co-transfecting them with 3^rd^ generation lentiviral packaging plasmids with TransIT-Lenti Transfection Reagent (Mirus cat. no. MIR6606) according to the manufacturer’s protocol. The next day, lentivirus was precipitated using Lentivirus Precipitation Solution (ALSTEM cat. no. VC150) according to the manufacturer’s protocol, resuspended in DPBS, and aliquoted and stored at -80 °C. iAstrocytes were transduced with lentivirus at the time of plating. When necessary, the functional titer of the lentivirus was determined by measuring BFP+ cells 48 hours after transducing iAstrocytes with a serial dilution of lentivirus. The protospacer sequences of the sgRNAs used in this study were selected from those generated in Horlbeck *et al.* (37) and are listed below (when two protospacers are listed, the sgRNA is cloned into pMK1334 using the Replogle *et al.* method):

Non-targeting controls (NTC):

GGAGTTAAGGCCTCGTCTAG, GTGCGGGGGCATGGCCCCGC

*MTOR*

GGGACAGCGGGGAAGGCGGG

*RPTOR*:

GACCCCAGGTCCCAAGCCAC, GTGGGGGCCGCTAGGAAATG

*RICTOR*:

GGCGCAGGGCGGAATGACAG, GAGCGGGCTTACCTCGTACT

*TSC1*:

GGGCCTTGGCCCTTTCACGA

*VAMP7*:

GGCCAACTGCCCGCTCCCAG, GAGGGACGCGGGTCAGTGCA

*VAMP3*:

GCGGCAGCGGCGACGAGAGA, GGAAGCGAAGTTGGGACCGG

*SNAP23*:

GAACTCGGACACCCCAACAC, GTAGGGTGCAGCGCCAGGTC

*RAB27A/B*:

GACCCGAGCCAGGCGGGGAC (*RAB27A*), GCCCATCCGGAGCCACAGGT (*RAB27B*)

*IL32*:

GGAGCTGGGTCATCTCAGGT, GCGAAGGTGAGGACCCTCTG

### Generation and lentiviral transduction of reporter constructs

To subclone the original FIRE-pHLy construct (24) (pLBR10, Ubc:FIRE-pHLy) to make pLBR11 (GfaABC1D:FIRE-pHLy), we excised the FIRE-pHLy cassette from pLBR10 and linearized the backbone vector (pKL20; pHIV gfaABC1D:mRuby:WPRE) with BamH1-HF (New England BioLabs, cat. no. R3136) and ClaI (New England BioLabs cat. no. R0197). We then ran these reactions on a 1% agarose gel, gel extracted the relevant fragments, and ligated them together with T4 DNA Ligase (New England BioLabs cat. no. M0202) per manufacturer’s protocol to generate pLBR11. Both pLBR11 (all FIRE-pHLy experiments), and GFP-LC3-RFP-LC3ΔG (reporter cassette from Addgene plasmid #84572) (25) reporters were transduced into iAstrocytes with lentivirus at low multiplicity of infection (MOI < 1) during plating to prevent high expression of exogenous lysosomal proteins.

### Immunofluorescence imaging

For immunofluorescence imaging experiments involving iAstrocytes, iAstrocytes plated in Greiner μClear 96-well plates (Greiner Bio-One cat. no. 655087) were fixed with 4% paraformaldehyde (diluted from a 16% solution in DPBS; Electron Microscopy Sciences cat. no. 15710) for 15 min at RT. After washing three times with DPBS, we blocked and permeabilized cells with DPBS (Milipore Sigma cat. no. D8537) + 3% BSA (Milipore Sigma cat. no. A9647) + 0.1% Triton X-100 (Millipore Sigma cat. no. X100) for 30 min at RT. Primary antibodies against LC3 (1:50; Cell Signaling Technology cat. no. 2775), LAMP1 (1:50; Abcam cat. no. ab25630), LAMP2 (1:500; Abcam cat. no. ab25631), IL-32 (1:250; ProteinTech cat. no. 11079-1-AP), or CD63 (1:1420; BioLegend cat. no. 353039) were added to blocking buffer and incubated with iAstrocytes overnight at 4 °C. Afterwards, the samples were washed with DPBS three times, incubated with pre-adsorbed secondary antibodies (1:500 goat anti-mouse IgG Alexa Fluor 488, 1:500 goat anti-rabbit IgG Alexa Fluor 555; Abcam cat. no. ab150117 and ab150086) for 1 hour at RT, washed three times with DPBS, incubated with 1 μg/mL Hoechst 33342 (ThermoFisher Scientific cat. no. H3570) for 10 minutes and then washed two additional times before imaging on an IN Cell Analyzer 6000, using a 60X 0.7 NA objective, 2×2 binning, 100-400 ms exposure, an aperture width of ∼1 Airy unit, and 9-16 fields per well, or on an ImageExpress Confocal HT.ai (Molecular Devices), using an APO 20x 0.95 NA water-immersion objective (Nikon), 89 North LDI lasers 405 nm, 475 nm, and 555 nm with emission filters 452/32 nm 520/428 nm and 598/25 nm for Hoechst 333342, AlexaFluor 488 and 555, respectively, 1x1 binning (325 x 325 nm pixel size), 100-400 ms exposure, 60 μm pinhole spinning disk, and 12 fields per well captured on a Zyla 4.5 sCMOS camera (Andor).

For immunofluorescence imaging experiments involving HIE samples, all human HIE tissue was collected with informed consent and in accordance with guidelines established by the UCSF Committee on Human Research (H11170-19113-07), as previously described (85). Immediately after procurement, all brains were immersed in PBS with 4% paraformaldehyde for 3 days. On day 3, the brain was cut in the coronal plane at the level of the mammillary body and immersed in fresh 4% paraformaldehyde and PBS for an additional 3 days. After fixation, all tissue samples were equilibrated in PBS with 30% sucrose for at least 2 days. After sucrose equilibration, tissue was placed into molds and embedded with OCT medium for 30 minutes at room temperature, followed by freezing in dry-ice-chilled ethanol. UCSF neuropathology staff performed brain dissection and its evaluation. The diagnosis of HIE requires clinical and pathological correlation; no widely accepted diagnostic criteria are present for the pathological diagnosis of HIE. HIE cases showed consistent evidence of diffuse white matter gliosis, as evaluated by the qualitative increase in the number of GFAP^+^ cells in addition to the increased intensity of GFAP staining. For immunostaining of reactive astrocyte markers, tissue slides were bleached with UV overnight, rinsed with PBS for 10 minutes and incubated with blocking solution (10% normal goat serum + 0.2% Triton X-100 in PBS) for 1 hour at room temperature and then incubated with the following primary antibodies overnight at 4 °C: chicken anti-GFAP (1:500 dilution; BioLegend cat. no. 829401), mouse anti-NeuN (1:500 dilution; Millipore Sigma cat. no. MAB377), mouse anti-OLIG1 (1:200; Millipore Sigma cat. no. MAB2417), or rabbit anti IL32 (1:200 dilution; ProteinTech cat. no. 11079-1-AP). Afterwards, the samples were rinsed three times with PBS + 0.2% Triton X-100 for 10 minutes each time and then incubated with the appropriate secondary antibodies for 1 hour at room temperature in the dark.

### LysoTracker staining and imaging

iAstrocytes were washed with DPBS and incubated with a 50 nM solution of LysoTracker Green DND-26 (ThermoFisher Scientific cat. no. L7526) or LysoTracker Red DND-99 (ThermoFisher Scientific cat. no. L7528) in DPBS for 5 minutes at 37°C, washed two times with DPBS, and processed for downstream experiments. For imaging, iAstrocytes were imaged at 37°C and 5% CO_2_ using an IN Cell Analyzer 6000 as above. For flow cytometry (CRISPRi screens), iAstrocytes were dissociated with Accutase for 10 mins at 37°C after LysoTracker staining, and resuspended in FACS buffer (DPBS + 1% BSA + 2 mM EDTA (Milipore Sigma cat. no. 324506)).

### CTSB activity assay

Intracellular CTSB activity was measured using the Cathepsin B Assay Kit (Magic Red) from Abcam (ab270772) following the manufacturer’s instructions. Vehicle vs ITC-treated iAstrocytes plated on a Matrigel-coated Greiner μClear 96-well plates (Greiner Bio-One cat. no. 655087) were pre-treated with vehicle or 100 nM Bafilomycin A1 for 3 hours and then incubated with 7 uL of 1:10 diluted Magic Red CTSB substrate per well for 1.5 hours. Fluorescence from cleaved Magic Red CTSB substrate was then measured on a SpectraMax M5 fluorescence plate reader with excitation set to 592 nm and emission set to 628 nm with auto-cutoff enabled. iAstrocytes were then incubated with 10 μg/mL Hoechst for 15 min at 37°C and then read on the plate reader with excitation set to 359 nm and emission set to 461 nm with auto-cutoff enabled. CTSB activity normalized to cell number was calculated by dividing the Magic Red fluorescence by the Hoechst fluorescence.

### Total internal reflection fluorescence microscopy

iAstrocytes were plated at 20,000 cells/cm2 in 35 mm imaging dishes (Ibidi, cat. no. 81156) and simultaneously transduced with pLBR11 (GfaABC1D:FIRE-pHLy). Prior to imaging, the conditioned media of iAstrocytes was collected. iAstrocytes were then stained with LysoTracker as detailed above and re-incubated with the collected conditioned media for the duration of imaging. TIRF was performed on a Nikon Ti-E inverted microscope equipped with a Nikon TIRF illuminator, Nikon motorized stage with piezo Z inset, Andor iXon Ultra DU897 camera, and run by NIS-Elements (v5.20.00 build 1423). iAstrocytes were imaged using an Apo TIRF 100x/1.49 oil (DIC N2/100X I) objective, 488nm and 561nm lasers (Agilent MLC400 monolithic laser combiner) and Sutter Lambda 10-B with ET525/50m and ET600/60m emission filters (Chroma), respectively, for 120 seconds with 100 ms exposure (per channel).

### Antibody staining for flow cytometry

To measure cell-surface protein levels (i.e. LAMP1, TFRC), iAstrocytes were dissociated with Accutase for 10 mins at 37 °C, diluted with ice-cold DPBS + 1% BSA, and spun down at 300 x g for 5 minutes at 4 °C. After aspirating the supernatant, all cells were incubated in 1:20 Fc block (BD Bioscience cat. no. 564220) diluted in DPBS + 3% BSA for 10 minutes on ice, followed by the addition of 1:50 primary antibodies against LAMP1 (AF488- or BV605-conjugated; BioLegend cat. no. 328610 and 328634) for an additional 30 minute incubation on ice in the dark. iAstrocytes were then washed with ice-cold DPBS + 1% BSA, spun down at 300 x g for 5 mins at 4 °C, and resuspended in DPBS + 1% BSA for flow cytometry. When astrocytes were not transduced with sgRNAs, DAPI (ThermoFisher Scientific cat. no. D1306) was added at 0.4 mg/mL to assess cell viability; when astrocytes were transduced with sgRNAs, TO-PRO-3 (ThermoFisher Scientific cat. no. T3605) was added at 1 μM.

For intracellular staining, dissociated iAstrocytes were fixed with 2% paraformaldehyde for 10 min at RT, washed twice with DPBS + 0.5% Tween 20 (Millipore Sigma cat. no. P9461), incubated with unconjugated primary antibodies against C3 (1:500; ThermoFisher Scientific cat. no. PA1-29715), VCAM1 (1:250; BioLegend cat. no. 305802), phospho-S6 (1:200; Cell Signaling Technology cat. no. 2211), phospho-4E-BP1 (1:200; Cell Signaling Technology cat. no. 2855), p62 (1:200; Cell Signaling Technology cat. no. 88588), or IL-32 (1:250; ProteinTech cat. no. 11079-1-AP) for 20 min at RT, washed with DPBS + 0.5% Tween 20, incubated with conjugated secondary antibodies (1:1000, Donkey anti mouse IgG AlexaFluor 488 conjugated, ThermoFisher Scientific cat. no. A-21202; 1:1000 Donkey anti rabbit IgG AlexaFluor 555 conjugated, ThermoFisher Scientific cat. no. A-31572; 1:1000 Donkey anti goat IgG AlexaFluor 647 conjugated, ThermoFisher Scientific cat. no. A-21447) for 20 min at RT, washed with DPBS + 0.5% Tween 20, and then resuspended in DPBS + 0.5% Tween 20 for flow cytometry.

### Pooled CRISPRi screening

CRISPRi screens were conducted as described in Leng *et al*. (9). Both LysoTracker and LAMP1 screens were conducted with the H1 (i.e. “druggable genome”) sgRNA sub-library from our next-generation CRISPRi library (37). We generated lentivirus for the pooled H1 sgRNA library as previously described (82). For each screen (i.e. LysoTracker or cell-surface LAMP1, vehicle or ITC), iAstrocytes were plated at 20,000 cells/cm^2^ onto 4 Matrigel-coated 15-cm dishes, transduced with the H1 lentiviral sgRNA library with >70% transduction efficiency, treated with vehicle control or IL-1α+TNF+C1q for 24 hours, stained for LysoTracker or cell-surface LAMP1, and then sorted into LysoTracker high vs. low or cell-surface LAMP1 high vs. low (top and bottom 35% of cells on fluorescence histogram) populations. iAstrocytes were sorted with a BD FACSAria Fusion cell sorter at 5,000-10,000 events per second, and then pelleted via centrifugation at 300 x g for 10 minutes for genomic DNA extraction. sgRNA abundances were then measured using next-generation sequencing as previously described (82). LysoTracker screens were performed with two experimental replicates per condition, while LAMP1 screens were done with one replicate. Results from all CRISPRi screens in this study can be interactively explored at www.crisprbrain.org (86).

### Extracellular vesicle isolation

For all experiments involving isolation of extracellular vesicles (EVs) from astrocyte conditioned media, astrocytes were treated with ScienCell Astrocyte media made using exosome-depleted FBS (ThermoFisher Scientific cat. no. A2720803). Extracellular vesicles were isolated from 10 mL of condition media from iAstrocytes plated onto 10 cm dishes using differential ultracentrifugation as described in Patel *et al.* (87). All spins were done at 4 °C. Briefly, conditioned media was spun down at 300 g for 10 min, then the supernatant was spun down at 2000 g for 30 min, then the supernatant was spun down at 15,600 g for 30 min, then the supernatant was spun down at 120,000 g for 2 hours, and finally the pellet (not visible) was resuspended in 100 μL of DPBS. The spins at 15,600 g and 120,000 g were done using a SW 41 Ti rotor with 13.2 mL UltraClear tubes (Beckman Coulter cat. no. 344059) in a Beckman Coulter Optima XE ultracentrifuge. To increase exosome yield for western blotting against certain targets shown in Fig. 5a (Hsc70, Caveolin-1, IL-32), 10 nM bafilomycin A1 was added together with vehicle control or ITC. For IL-32, bands were present on blots using samples treated with or without bafilomycin A1 (see raw blot image files in Supplementary File 1).

### Measurement of EV size distribution

The size distribution of EVs isolated above was measured via nanoparticle tracking analysis using the NanoSight 3000 following the manufacturer’s instructions. 100 μL of each resuspended EV sample was diluted to final volume of 500 μL for analysis on the Nanosight 3000.

### Western blots

iAstrocytes were dissociated with Accutase for 10 mins at 37°C, spun down at 300 x g for 5 mins, washed with ice-cold DPBS, lysed in RIPA buffer (Thermo Fisher Scientific, cat. no. 89900) with protease (Roche, cat. no. 04693159001) and phosphatase inhibitors (Sigma-Aldrich, cat. no. PHOSS-RO), and incubated on ice for 45 minutes with brief vortexing every 15 minutes. Samples were centrifuged at 18,000 x g for 20 mins at 4°C to pellet insoluble components of lysate. After the total protein was quantified using the Pierce BCA Protein Assay Kit (ThermoFisher Scientific, cat. no. 23225) in the supernatant, each sample was diluted to the same concentration, and denatured with NuPAGE LDS Sample Buffer (4X) (ThermoFisher Scientific, cat. no. NP0007) and reduced with DTT.

Equivalent protein amounts were run on 10% NuPAGE Bis-Tris gels (ThermoFisher Scientific, cat. no. NP0301) and transferred to nitrocellulose membranes (Bio-Rad, 1704271). For blots shown in Fig. 1f-h, Fig. 1l, and Fig. 3g, 20-40 μg of protein was loaded; for blots shown in Fig. 5a,c, 1-2 μg of protein was loaded. We performed ponceau staining (Milipore Sigma, cat. no. P7170) to validate equivalent loading between lanes. Membranes were blocked with Intercept blocking buffer (LI-COR cat. no. 927-60001) on a shaker, after which they were incubated with primary antibodies against mTOR (1:1000, Cell Signaling Technology cat. no. 2972), phospho-S6 (1:500; Cell Signaling Technology cat. no. 2211), S6 (1:1000; Cell Signaling Technology cat. no. 2217), phospho-ULK1 (1:500; Cell Signaling Technology cat. no. 14202), phospho-Akt (1:500; Cell Signaling Technology cat. no. 4060), pan-Akt (1:1000; Cell Signaling Technology cat. 2920), phospho-4E-BP1 (1:1000; Cell Signaling Technology cat. no. 2855), LAMP1 (1:1000; Abcam cat. no. ab25630), LAMP2 (1:1000; Abcam cat. no. ab25631), CTSD (1 ug/mL, R&D Systems cat. no. AF1014), β-actin (1:1000; Cell Signaling Technology cat. no. 4970), EGFP (1:200; Santa Cruz Biotechnology cat. no. sc-9996), LC3 (1:1000; Cell Signaling Technology cat. no. 2775), CD81 (1:500; BioLegend cat. no. 349502), CD63 (1:5680; BioLegend cat. no. 353039), Apo-AI (1:200; Santa Cruz Biotechnology cat. no. sc-376818), GAPDH (1:200; Santa Cruz Biotechnology cat. no. sc-47724), Hsc70 (1:200; Santa Cruz Biotechnology cat. no. sc-7298), TSG101 (1:500; BioLegend cat. no. 934301), ALIX (1:500; BioLegend cat. no. 634501), or Caveolin-1 (1:1000; Cell Signaling Technology cat. no. 3267) at 4°C overnight on a shaker. Blots were washed with TBS and incubated with secondary antibodies goat anti-mouse IRDye 800CW (1:10,000; LI-COR cat. no. 926-32210) and goat anti-rabbit IRDye 680RD (1:10,000; LI-COR cat. no. 926-68071) in blocking buffer (see above) for 1 hour at RT on a shaker. Afterwards, blots were washed with TBS, and imaged using an Odyssey Fc Imaging system (LI-COR, cat. no. 2800). To re-probe blots, we incubated membranes with 1X NewBlot Nitro Stripping Buffer (LI-COR, cat. no. 928-40030) for 5 minutes at RT on a shaker before re-blocking and probing as above. Immunoblot bands were quantified with LI-COR ImageStudio software. For blotting against CD63 and CD81, reducing agent was omitted from the sample buffer since the antibodies used here recognize epitopes containing disulfide bonds.

Source data for immunoblots is provided in Supplementary File 1.

### mito-EV staining for flow cytometry

Conditioned media from iAstrocytes was spun down at 300 g to remove dead cells and then incubated with 10 μg/mL Hoechst, 5 nM TMRM (ThermoFisher Scientific cat. no. T668), and 50 nM MitoTracker Green (ThermoFisher Scientific cat. no. M7514) for 5 min at 37 °C. After incubation, the conditioned media was analyzed directly with flow cytometry.

### IL-32 or CTSB immunoassays

IL-32 concentration was measured using an ELISA kit (R&D Systems cat no. DY3040-05) following the manufacturer’s instructions. For measurement of IL-32 concentration in iAstrocyte conditioned media, the conditioned media was first spun down at 300 g to remove dead cells, then 50 μL of conditioned media + 25 μL of RIPA buffer + 25 μL of 1% BSA in DPBS was added per well, with the same fraction of RIPA and diluent added to the IL32 standards. For measurement of IL-32 concentration in the exosome fraction vs. supernatant, 75 μL of sample + 25 μL of RIPA buffer was added per well, with the same fraction of RIPA and diluent added to the IL32 standards. CTSB concentration was measured via an electrochemiluminescence immunoassay (Mesoscale Discoveries cat. no. K151ACAR-2) following the manufacturer’s instructions.

### Whole-cell proteomics sample preparation

iAstrocytes were washed and collected (detached gently using a cell scraper) in ice-cold DPBS, spun down at 300 x g for 5 mins at 4 °C, and stored at -80 °C. Cell pellets were lysed in 2.5% sodium dodecyl sulfate (SDS) 50mM Tris pH 8 and heated to 95 °C for 5min, then subjected to probe sonication. Lysates were centrifuged at 14000g for 15 min before protein quantification using BCA assay (Pierce, Thermo Fisher Scientific) according to manufacturer’s instructions. Protein lysates were acetone precipitated by adding 5 volumes of ice-cold acetone, vortexed, incubated at -20 °C for 1 hr and pelleted by centrifugation at 21000g and 4 °C for 15 min.

Protein pellets were resuspended in 1% sodium deoxycholate (SDC) 50 mM EPPS pH 8.5 and reduced by incubating at 37 °C for 20 min with rotation in the presence of 5mM DTT. Protein was then alkylated by incubating in the dark at 37 °C for 20 min in the presence of 25mM iodoacetamide before digestion overnight with LysC at an enzyme-to-protein ratio of 1:50 followed by 3 hr digestion with trypsin at an enzyme-to-protein ratio of 1:50. Digestion was stopped by acidification to 1% trifluoroacetic acid (TFA) and placed on ice for 10min to promote precipitation of SDC. The SDC was then pelleted and removed by centrifugation at 21000g for 10min. The supernatant containing acidified peptides were then de-salted on in-house made Stagetips (88) packed with polystyrene divinyl benzene reverse phase sulphonate solid phase extraction material (CDS Empore, Fisher Scientific) as previously published (88). Stagetips were prepared by activating with 100% methanol, conditioning with 80% acetonitrile containing 0.1% TFA, equilibrated with 0.2% TFA, followed by sample loading, washing twice with 99% isopropanol containing 0.1% TFA, washing twice with 0.2% TFA and once with 0.1% FA. Peptides were eluted with 60% acetonitrile (ACN) containing 0.5% ammonium hydroxide. The desalted peptides were then flash frozen and dried by centrifugal evaporation.

### Endolysosomal proteomics sample preparation

pLBR09 (gfaABC1D: LAMP1-XTEN80-mEGFP-3XHA) used for lysosome immunoprecipitation was generated by subcloning pLBR08 (EF1α: LAMP1-XTEN80-mEGFP-3XHA). Briefly, pLBR08 was generated via Gibson assembly of PCR-amplified LAMP1 cDNA (from mTagRFP-T-Lysosomes-20 acquired from Nikon Imaging Center at UCSF, Addgene plasmid #58022) and PCR-amplified XTEN80-mEGFP-3xHA immunoprecipitation tag with a linearized backbone generated from ClaI and BspDI (New England BioLabs cat. no. R0557) digestion of pKL017 (pHIV EF1a:Clover:WPRE). pLBR09 was then generated by linearizing pKL020 (pHIV gfaABC1D:mRuby:WPRE) and excising the LAMP1-XTEN80-mEGFP-3XHA cassette of pLBR08 via restriction digest with XbaI (New England Biolabs cat. no. R0145) and ClaI. The resulting fragments were run on a 1% agarose gel and gel extracted (Zymoclean Gel DNA Recovery Kit, cat. no. D4007). These fragments were ligated together with T4 DNA Ligase per manufacturer protocol. iAstrocytes were plated at 20,000 cells/cm^2^ per experimental replicate and transduced with pLBR09 lentivirus during plating. We used three replicates each for vehicle and ITC conditions.

Lysosome immunoprecipitation was performed as previously described (89). Briefly, iAstrocytes were washed and collected (detached gently using a cell scraper) in ice-cold DPBS, spun down at 300 x g for 5 mins at 4 °C. We then resuspended the cell pellet in homogenization buffer (25 mM Tris HCl pH 7.5, 50 mM sucrose, 0.5 mM MgCl_2_, 0.2 mM EGTA with protease inhibitors). The iAstrocyte cell suspension was mechanically lysed with a 23G syringe, immediately transferred to isotonic buffer (2.5M sucrose, 0.2 mM EGTA, 0.5 mM MgCl_2_) and spun at 3000 x g for 10 mins at 4 °C. The supernatant was transferred to anti-HA magnetic beads (Thermo Scientific cat. no. PI88836) for the pulldown of intact lysosomes, with the final resuspension in 1X urea buffer (2M Urea, 1 mM DTT, 12.5 mM Tris). On-bead digest was performed by sequentially incubating the resulting protein samples with 5 mM (final concentration) IAA at 22°C and 225 rpm for 30 mins, 0.5 μg LysC at 22°C and 225 rpm overnight, 1 μg of trypsin at 22°C and 225 rpm for 4 hours, and 1% (final concentration) TFA. Acidified peptides were then de-salted according to the same protocol as whole-cell proteome samples. For western blotting, NuPAGE LDS Sample Buffer (4X) was added to samples collected prior to the on-bead digestion. See Western blots section for further details.

### LC-MS Analysis

Peptides were resuspended in 2% ACN with 0.1% TFA before loading onto a 25 cm x 75 µm ID, 1.6 µm C18 column (IonOpticks) maintained at 40°C. Peptides were separated with an EASY-nLC 1200 system (Thermo Fisher Scientific, San Jose, CA) at a flow rate of 300 nL min^-1^ using a binary buffer system of 0.1% FA (buffer A) and 80% acetonitrile with 0.1% FA (buffer B) in a two-step gradient. The following was performed for whole cell proteome samples, 3% to 27% B in 105 min and from 27% to 40% B in 15min. For lysosome-specific samples, 3% to 27% B in 52.5 min and from 27% to 40% B in 14.5 min. All samples were analyzed on a Fusion Lumos mass spectrometer (Thermo Fisher Scientific, San Jose, CA) equipped with a nanoFlex ESI source operated at 1550 volts, RF lens set to 30%, operated in data dependent acquisition mode with a duty cycle time of 1 sec. Full MS scans were acquired with a m/z scan range of 375-1500 m/z in the Orbitrap mass analyzer (FTMS) with a resolution of 240k for whole cell proteome and 120k for lysosome-specific samples. Selected precursor ions were subjected to fragmentation using higher-energy collisional dissociation (HCD) with a quadrupole isolation window of 0.7 m/z, and normalized collision energy of 31%. HCD fragments were analyzed in the Ion Trap mass analyzer (ITMS) set to Turbo scan rate. Fragmented ions were dynamically excluded from further selection for a period of 45 sec for whole cell proteome and 60 sec for lysosome-specific samples. The AGC target was set to 1,000,000 and 10,000 for full FTMS and ITMS scans, respectively. The maximum injection time was set to Auto for both full FTMS and ITMS scans.

### Drug treatments

Vacuolin-1 (Milipore Sigma, cat. no. 673000), bafilomycin A1 (Milipore Sigma, cat. no. B1793), rapamycin (Milipore Sigma, cat. no. 553210), PP242 (Milipore Sigma, cat. no. 475988), RapaLink-1 (MedChemExpress cat. no. HY-111373), PI4KIII beta inhibitor 3 (MedChemExpress cat. no. HY-15679), and PI-273 (MedChemExpress cat. no. HY-103489) were resuspended in DMSO per manufacturer’s instructions, aliquoted, and stored at -80°C. All compounds were added to iAstrocytes alongside vehicle or ITC treatments (i.e. 24 hour treatments) with the exception of using bafilomycin A1 for measurement of autophagic flux, which was added to astrocytes for 4 hours before sample collection.

### Data analysis

#### Analysis of published RNA-seq datasets

We used BioJupies (90) to reanalyze bulk RNA-seq data and obtain differentially expressed genes from Perriot *et al*. (91) (GSE120411; hiPSC-derived astrocytes treated with IL1β and TNF) and Barbar *et al*. (19) (syn21861229; CD49f+ astrocytes sorted from cerebral organoids treated with vehicle control or IL-1α+TNF+C1q).

#### Overlap analysis of differentially expressed genes (DEGs)

Human or rodent astrocyte DEGs from various diseases or disease models were extracted manually from the supplementary information provided on the respective publications or shared by the corresponding authors upon request; see Supplementary Table 3 for a list of all publications and datasets used for the analysis. GO Cellular Component terms relating to the endolysosomal system and biological pathway terms related to mTOR signaling were downloaded from Enrichr (92–94). Overlap between astrocyte DEGs and the gene sets of interest was assessed and visualized using the R package GeneOverlap (v1.38.0); Fisher’s exact test *P* values were adjusted for multiple testing using the Benjamini-Hochberg procedure.

#### Proteomics

Raw files were searched against the human reference proteome (Swiss-Prot, downloaded on 20^th^ of April 2020, refer to PRIDE upload for full sequence) along with the default common contaminants list with enzyme specificity set to trypsin within MaxQuant version 2.0.1.0 using standard settings with match between runs and the MaxLFQ algorithm activated. Whole cell proteome (raw data file names begin with ‘Proteome’) and lysosome-specific (raw data file names begin with ‘LAMP1’) raw files were set as separate parameter groups to ensure that the MaxLFQ algorithm was performed only within each sample set. The resulting proteinGroups text file was then uploaded to Perseus version 1.6.7.0 where all reverse sequences, potential contaminants and proteins only identified by site were filtered out of the dataset. All protein abundance values were normalized by Log 2 transformation. The lysosome-specific raw data files were evaluated separate from whole cell proteomic files. Search results from lysosomal raw data files were filtered to include proteins that were quantified in at least two out of three biological replicates within at least the ITC (raw data file names include ITC) or vehicle (raw data file names include WT) triplicate set. Missing values were then imputed within each replicate by replacing from the normal distribution using default settings. A two sample students t-test was performed between the ITC treated and vehicle triplicates, with permutation-based FDR using default settings. The resulting T-test difference and significance were used to generate the lysosome-specific volcano plot.

#### Pathway enrichment analysis

We used Enrichr (92–94) to perform enrichment analysis of gene lists.

#### CRISPRi screening analysis

We analyzed next generation sequencing reads generated from our CRISPRi screens as previously described (82). The “gene score” was calculated as -log_10_(*P* value) times the phenotype score.

#### Pseudobulk analysis of snRNA-seq data from Macnair et al

The processed feature-barcode count matrix from Macnair *et al.* (62) was downloaded from https://zenodo.org/records/8338963 (95) and loaded into R as a Seurat object.

Seurat::AggregateExpression was then used to combine raw counts across cells from each cell type for each sample. DESeq2::estimateSizeFactors was then used to normalize the counts, and then the normalized counts added a pseudocount of 1 followed by transformation to log_2_ scale. IL-32 log-normalized counts were then plotted across samples derived from different multiple sclerosis lesions. *P* values for differences in IL-32 log-normalized counts were calculated by linear regression in R using lm() without adjustment for clinical covariates.

#### Flow cytometry

Data from flow cytometry experiments were analyzed using FlowJo (version 10.7.1). Live cells were gated by plotting SSC-A vs. FSC-A and then single cells were gated by plotting FSC-H vs. FSC-A. For experiments involving CRISPRi knockdown, analysis was restricted to sgRNA-transduced cells (gating on the histogram of BFP fluorescence values). For antibody staining experiments where median fluorescence intensity (MFI) values were reported, the average MFI of unstained control samples were subtracted from the MFI of stained samples.

#### Fluorescence imaging analysis

We used CellProfiler (v3.15) (96) to quantify iAstrocyte immunostaining imaging data. The total image intensity of LAMP1, LAMP2, LC3, and LysoTracker signals were quantified after first thresholding images relative to either a no-primary antibody control (immunofluorescence— LAMP1, LAMP2, LC3) or no-dye control (LysoTracker). The total image intensity of LAMP1, LAMP2, and LC3 was then divided by the total intensity of Hoechst to correct for variation in cell number. All frames collected were averaged to return a single quantification per well, which we considered as a single data point for statistical tests.

#### Statistics and reproducibility

Sample sizes were determined by referencing existing studies in the field. Major findings were validated using independent samples and orthogonal approaches. Numbers of replicates are listed in each figure. Randomization was not relevant to our study because no animals or human subjects were involved. Statistics were computed in RStudio (version 4.0.5). Normality of data distributions were tested where appropriate (Shapiro-Wilk test using the function shapiro.test).

Simple pairwise tests were conducted using t.test for normally distributed data or wilcox.test for non-normally distributed data. For instances involving multiple comparisons of normally distributed data, linear regression using the function lm was used to compute *P* values. For experiments with factorial design, linear regression including interaction terms (equivalent to multi-way ANOVA) was performed using the function lm. For percentage data or data ranged from 0 to 1, beta regression was used to calculated *P* values using the R package betareg (v3.1-4), with logit linking function and the bias-corrected maximum likelihood estimator. Correction for multiple testing was performed where appropriate.

## LIST OF ABBREVIATIONS

CRISPR: clustered regularly interspaced short palindromic repeats
CRISPRi: CRISPR-based gene expression inhibition
hiPSC: human induced pluripotent stem cell
ITC: IL-1α+TNF+C1q
ELISA: enzyme-linked immunosorbent assay
EV: extracellular vesicle
EAE: experimental autoimmune encephalitis

