## Supplementary File 1 for "mTOR activation induces endolysosomal remodeling and nonclassical secretion of IL-32 via exosomes in inflammatory reactive astrocytes": BRExp225h2_metadata.docx

| Sample # | CytokineTreatment | DrugTreatment | Blot # |
| --- | --- | --- | --- |
| 1 | Veh | 0.1% DMSO | 1 |
| 2 | Veh | 0.1% DMSO | 1 |
| 3 | Veh | 0.1% DMSO | 1 |
| 4 | ITC | 0.1% DMSO | 1 |
| 5 | ITC | 0.1% DMSO | 1 |
| 6 | ITC | 0.1% DMSO | 1 |
| 7 | Veh | 100 nM Bafilomycin A | 1 |
| 8 | Veh | 100 nM Bafilomycin A | 1 |
| 9 | Veh | 100 nM Bafilomycin A | 1 |
| 10 | ITC | 100 nM Bafilomycin A | 1 |
| 11 | ITC | 100 nM Bafilomycin A | 1 |
| 12 | ITC | 100 nM Bafilomycin A | 1 |
| 13 | Veh | 0.1% DMSO | 2 |
| 14 | Veh | 0.1% DMSO | 2 |
| 15 | Veh | 0.1% DMSO | 2 |
| 16 | ITC | 0.1% DMSO | 2 |
| 17 | ITC | 0.1% DMSO | 2 |
| 18 | ITC | 0.1% DMSO | 2 |
| 19 | Veh | 100 nM Bafilomycin A | 2 |
| 20 | Veh | 100 nM Bafilomycin A | 2 |
| 21 | Veh | 100 nM Bafilomycin A | 2 |
| 22 | ITC | 100 nM Bafilomycin A | 2 |
| 23 | ITC | 100 nM Bafilomycin A | 2 |
| 24 | ITC | 100 nM Bafilomycin A | 2 |

Round 1

Blot 1 & 2, top (>37 kDa):

Rb anti p-ULK1 (CST 6888), 1:1000 -> Gt anti Rb IRDye680 1:10,000

Rb anti p-Akt (CST 9271), 1:1000 -> Gt anti Rb IRDye680 1:10,000

Ms anti SQSTM1/p62 (CST 88588), 1:1000 -> Gt anti Ms IRDye800 1:10,000

Blot 1 & 2, bot (<37 kDa):

Rb anti LC3B (CST 2775), 1:1000 -> Gt anti Rb HRP 1:10,000

Ms anti GAPDH (sc-47724), 1:500 -> Gt anti Ms IRDye800 1:10,000
