## Supplementary File 1 for "mTOR activation induces endolysosomal remodeling and nonclassical secretion of IL-32 via exosomes in inflammatory reactive astrocytes": mTOR_GAPDH_pULK1_metadata.docx

**Last edited by BR 03/23/24 12pm EST.**

**Exp220:**

| **Metadata for BR western blots** | | | | | |
| --- | --- | --- | --- | --- | --- |
| Experiment | Replicate | Raw Image Name | Channel | Replicate |  |
| Exp220c | 1 | 0001379_01 | 700 | 2 | MTOR |
| Exp220c | 1 | 0001380_01 | 700 | 1 | MTOR |
| Exp220c | 1 | 0001381_01 | 700 | 3 | MTOR |
| Exp220c | 1 | 0001379_01 | 800 | 2 | GAPDH |
| Exp220c | 1 | 0001380_01 | 800 | 1 | GAPDH |
| Exp220c | 1 | 0001381_01 | 800 | 3 | GAPDH |
| Exp220c | 2 | 0001392_01 | 800 | 1 | p-ULK1 (S757) |
| Exp220c | 2 | 0001393_01 | 800 | 2 | p-ULK1 (S757) |
| Exp220c | 2 | 0001394_01 | 800 | 3 | p-ULK1 (S757) |
| Exp220c | 2 | 0001562_01 | 800 | 1 | p-AKT (S473) |
| Exp220c | 2 | 0001563_01 | 800 | 2 | p-AKT (S473) |
| Exp220c | 2 | 0001564_01 | 800 | 3 | p-AKT (S473) |
| Exp220c | 2 | 0001571_01 | 800 | 1 | AKT |
| Exp220c | 2 | 0001570_01 | 800 | 2 | AKT |
| Exp220c | 2 | 0001569_01 | 800 | 3 | AKT |
| Exp220f | 1 | 0001663_01 | 800 | 1 | p-S6 |
| Exp220f | 1 | 0001664_01 | 800 | 1 | p-S6 |
| Exp220f | 1 | 0001665_01 | 800 | 1 | p-S6 |
| Exp220f | 1 | 0001691_01 | 800 | 2 | S6 |
| Exp220f | 1 | 0001692_01 | 800 | 2 | S6 |
| Exp220f | 1 | 0001693_01 | 800 | 2 | S6 |

**See below for experimental details.**

**Exp220c**

R1: total MTOR and GAPDH blots in manuscript (Fig. 3g).

Antibodies

- mTOR (1:1000, polyclonal rabbit; Cell Signaling Technology cat. No. 2972S)
  - Goat anti-rabbit IRDye 680RD (1:10,000; LI-COR cat. no. 926-68071)
- phospho-ULK1 (1:500; polyclonal rabbit; Cell Signaling Technology cat. no. 14202)
  - Goat anti-rabbit IRDye 680RD (1:10,000; LI-COR cat. no. 926-68071)
- GAPDH (1:200; Santa Cruz Biotechnology cat. no. sc-47724)
  - Goat anti-mouse IRDye 800CW (1:10,000; LI-COR cat. no. 926-32210)

R2: p-ULK1, p-AKT, and total AKT blots in manuscript (Fig. 3g, Fig. 3i).

Antibodies

- phospho-ULK1 (1:500; polyclonal rabbit; Cell Signaling Technology cat. no. 14202)
  - Goat anti-rabbit IRDye 680RD (1:10,000; LI-COR cat. no. 926-68071)
- phospho-AKT (1:500; monoclonal rabbit; Cell Signaling Technology cat. no. 4060)
  - Goat anti-rabbit IRDye 680RD (1:10,000; LI-COR cat. no. 926-68071)
- pan-Akt (1:1000; monoclonal mouse; Cell Signaling Technology cat. 2920)
  - Goat anti-mouse IRDye 800CW (1:10,000; LI-COR cat. no. 926-32210)

**Exp220f**

R1: p-S6 and S6 blots in manuscript (Fig. 3g).

Antibodies

- phospho-S6 (1:500; polyclonal rabbit; Cell Signaling Technology cat. no. 2211)
  - Goat anti-rabbit IRDye 680RD (1:10,000; LI-COR cat. no. 926-68071)
- S6 (1:1000; monoclonal rabbit; Cell Signaling Technology cat. no. 2217)
  - Goat anti-rabbit IRDye 680RD (1:10,000; LI-COR cat. no. 926-68071)
