## Supplementary File 1 for "mTOR activation induces endolysosomal remodeling and nonclassical secretion of IL-32 via exosomes in inflammatory reactive astrocytes": BRExp225g_metadata.docx

Samples 1-6: Veh

Samples 7-12: ITC

Blot 1:

Lanes 2-13: samples 1-12

Blot 2:

Lanes 3-14: samples 1-12

Round 1

Blot 1, top (>37 kDa):

Rb anti p-ULK1 (CST 6888), 1:1000 -> Gt anti Rb IRDye680 1:10,000

Ms anti LAMP1 (Abcam ab25630), 1:1000 -> Gt anti Ms IRDye800 1:10,000

Blot 1, bot (<37 kDa):

Rb anti p-S6 (CST 4858), 1:2000 -> Gt anti Rb IRDye680 1:10,000

Ms anti GAPDH (sc-47724), 1:500 -> Gt anti Ms IRDye800 1:10,000

Blot 2, top (>37 kDa):

Rb anti p-Akt (CST 9271), 1:1000 -> Gt anti Rb IRDye680 1:10,000

Ms anti pan-Akt (CST 2920), 1:2000 -> Gt anti Ms IRDye800 1:10,000

Blot 2 bot (<37 kDa):

None

Round 2

Blot 1, top (>37 kDa):

Rb anti p-ULK1 (CST 14202), 1:1000 -> Gt anti Rb IRDye680 1:10,000

Blot 1, bot (<37 kDa):

Rb anti S6 (CST 2217), 1:1000 -> Gt anti Rb IRDye680 1:10,000

Blot 2, top (>37 kDa):

Rb anti p-Akt (9271), 1:1000 -> Gt anti Rb IRDye680 1:10,000

Blot 2 bot (<37 kDa):

Ms anti GAPDH (sc-47724), 1:500 Gt anti Ms IRDye800 1:10,000
