## Supplementary File 1 for "mTOR activation induces endolysosomal remodeling and nonclassical secretion of IL-32 via exosomes in inflammatory reactive astrocytes": Exp247_WB_metadata.docx

**Exp247b:**

Set #2:

Primary antibodies

- R&D Gt anti IL32 polyclonal (AF3040) 0.2 mg/mL; manufacturer recommended WB concentration 0.5 ug/mL
- ProteinTech Rb anti IL32 polyclonal (11079-1-AP) 0.45 mg/mL
- BioLegend Ms anti CD81 (349502) 0.5 mg/mL
- BioLegend Ms anti CD63 (353039) 2.84 mg/mL
- SCBT Ms anti Hsp70/Hsc70 (sc-137239) 200 ug/mL
- SCBT Ms anti GAPDH (sc-47724) 200 ug/mL
- CST Rb anti OMA1 (95473)
- CST Rb anti OPA1 (67589)

Blot #1 (non-reducing):

Round 1

Ms anti CD81 1:1000 (0.5 ug/mL) -> Hs anti Ms HRP 1:10,000 [correct band present]

Ms anti CD63 1:2840 (1 ug/mL) -> Hs anti Ms HRP 1:10,000 [correct band present]

Strip

Round 2

Ms anti GAPDH 1:200 -> Gt anti Ms IRDye680 (red) 1:10,000 [correct band present]

Rb anti OPA1 1:1000 -> Gt anti Rb IRDye800 (green) 1:10,000

Blot #2 (reducing):

Round 1

Gt anti IL32 1:400 (0.5 ug/mL) -> Dk anti Gt IRDye680 (red) 1:10,000 [correct band present]

Rb anti IL32 1:900 (0.5 ug/mL) -> Dk anti Rb IRDye800 (green) 1:10,000 [correct band present]

Strip

Round 2

Ms anti Hsp70/Hsc70 1:200 (1 ug/mL) -> Gt anti Ms IRDye680 (red) 1:10,000 [correct band present]

Rb anti OMA1 1:1000 -> Gt anti Rb IRDye800 (green) 1:10,000 [correct band present]

Set #1:

Primary antibodies

- BioLegend Ms anti ALIX (634501) 0.5 mg/mL
- BioLegend Rt anti TSG101 (934301) 0.5 mg/mL
- SCBT Ms anti apoA-I (sc-376818) 0.2 mg/mL
- BioLegend Ms anti CD81 (349502) 0.5 mg/mL
- BioLegend Ms anti CD63 (353039) 2.84 mg/mL

Blot #1 (non-reducing):

Round 1

Ms anti CD81 1:1000 (1 ug/mL) -> Gt anti Ms IRDye680RD (red) 1:10,000 [correct bands detected]

Ms anti CD63 1:2840 (1 ug/mL) -> Gt anti Ms IRDye680RD (red) 1:10,000 [correct bands detected]

Blot #2 (reducing):

Round 1

Ms anti ALIX 1:500 (1 ug/mL) -> Gt anti Ms IRDye800CW (green) 1:10,000 **no bands seen**

Ms anti apoA-I 1:200 (1 ug/mL) -> Gt anti Ms IRDye800CW (green) 1:10,000 **no bands seen**

Rt anti TSG101 1:500 (1 ug/mL) -> Gt anti Rt IRDye680LT (red) 1:10,000 [correct bands detected]

**Exp247c:**

Set 1:

Primary antibodies

- BioLegend Ms anti ALIX (634501) 0.5 mg/mL
- BioLegend Rt anti TSG101 (934301) 0.5 mg/mL
- BioLegend Ms anti TFRC FITC conjugated (334103) 0.1 mg/mL
- CST Rb anti Caveolin-1 (3267) 0.691 mg/mL
- SCBT Ms anti HSC70 (sc-7298) 0.2 mg/mL
- CST Rb anti actin (4970)

Blot #1 (reducing)

Round 1

> 70 kDa piece:

Ms anti ALIX 1:500 (1 ug/mL) -> Hs anti Ms HRP 1:10,000 [correct bands possibly present]

Ms anti TFRC FITC conjugated 1:200 (0.5 ug/mL) -> Hs anti Ms HRP 1:10,000 [correct bands present]

< 70 kDa piece:

Rt anti TSG101 1:500 (1 ug/mL) -> Gt anti Rt HRP 1:10,000 **no bands seen**

Air dried

Blot #2 (reducing)

Round 1

Rb anti Caveolin-1 1:1000 (0.7 ug/mL) -> Gt anti Rb IRDye800 (green) 1:10,000 [correct bands present]

Ms anti Hsc70 1:200 (1 ug/mL) -> Gt anti Ms IRDye680 (red) 1:10,000 [correct bands present]

Strip with nitro stripping buffer

Round 2

>37 kDa piece:

Rt anti TSG101 1:500 (1 ug/mL) -> Gt anti Rt IRDye680 (red) 1:10,000 [correct bands present]

Rb anti actin 1:1000 -> Gt anti Rb IRDye800 (green) 1:10,000 [correct bands present]

<37 kDa piece:

Gt anti IL32 1:400 (0.5 ug/mL) -> Dk anti Gt IRDye680 (red) 1:10,000 [correct bands present]
