## Supplementary File 1 for "mTOR activation induces endolysosomal remodeling and nonclassical secretion of IL-32 via exosomes in inflammatory reactive astrocytes": Exp247c_set2_metadata.docx

Primary antibodies

BioLegend Ms anti ALIX (634501) 0.5 mg/mL; ~95 kDa

BioLegend Rt anti TSG101 (934301) 0.5 mg/mL; ~50 kDa

CST Rb anti Caveolin-1 (3267) 0.691 mg/mL; ~21-24 kDa

SCBT Ms anti HSC70 (sc-7298) 0.2 mg/mL; ~70 kDa

Dako Rb anti GFAP (Z0334) 3.2 mg/mL

CST Rb anti actin (4970); ~45 kDa

R&D Gt anti IL32 polyclonal (AF3040) 0.2 mg/mL

Blot #2 (reducing)

Lanes 1-2: ladder

Lanes 3-8: EV pellets samples 1-6

Lanes 9-14: supernatant samples 1-6

Round 1

Ms anti ALIX 1:1000 (0.5 ug/mL) -> Gt anti Ms IRDye680 (red) 1:10,000

Ms anti Hsc70 1:400 (0.5 ug/mL) -> Gt anti Ms IRDye680 (red) 1:10,000

Rb anti actin 1:1000 -> Gt anti Rb IRDye800 (green) 1:10,000

Rb anti Caveolin-1 1:1000 (0.7 ug/mL) -> Gt anti Rb IRDye800 (green) 1:10,000

Blot #1 (reducing)

Lane 1: ladder

Lanes 2-7: EV pellets samples 1-6

Lane 8: ladder

Lanes 9-14: supernatant samples 1-6

Lane 15: ladder

Round 1

> 37 kDa piece:

Rt anti TSG101 1:1000 (0.5 ug/mL) -> Gt anti Rt IRDye680LT (red) 1:10,000

Rb anti GFAP 1:3200 (1 ug/mL) -> Gt anti Rb IRDye6800 (green) 1:10,000

< 37 kDa piece:

Gt anti IL32 1:400 (0.5 ug/mL) -> Dk anti Gt IRDye680 (red) 1:10,000

Rb anti Caveolin-1 1:1000 (0.7 ug/mL) -> Dk anti Rb IRDye800 (green) 1:10,000
