## Supplementary File 1 for "mTOR activation induces endolysosomal remodeling and nonclassical secretion of IL-32 via exosomes in inflammatory reactive astrocytes": Exp250a_WB_cellLysates_set1_metadata.docx

Set 1:

Primary antibodies:

R&D Gt anti IL32 polyclonal (AF3040) 0.2 mg/mL; ~20-25 kDa

CST Rb anti actin (4970); ~45 kDa

CST Rb anti p-STAT3 (9145); ~79-86 kDa

CST Ms anti STAT3 (9139); ~79-86 kDa

CST Rb anti p-STAT1 (9167); 84-91 kDa

CST Rb anti STAT1 (9172);

Blot 1

Lane 1: ladder

Lanes 2-7: samples 1-6

Lane 8: ladder

Lanes 9-14: samples 7-12

Lane 15: ladder

Round 1

Gt anti IL32 1:400 (0.5 ug/mL) -> Dk anti Gt IRDye680 1:10,000

Rb anti actin 1:2,000 -> Dk anti Rb IRDye800 1:10,000

Rb anti p-STAT1 1:1,000 -> Dk anti Rb IRDye800 1:10,000

strip w/ Nitro stripping buffer, air dry

Blot 2

Lanes 1-2: ladder

Lanes 3-14: samples 1-12

Lane 15: ladder

Round 1

Ms anti STAT3 1:1,000 -> Gt anti Ms IRDye680 1:10,000

Rb anti p-STAT3 1:1,000 -> Gt anti Rb IRdye800 1:10,000

strip w/ Nitro stripping buffer, air dry
