## Supplementary figures and images for "mTOR activation induces endolysosomal remodeling and nonclassical secretion of IL-32 via exosomes in inflammatory reactive astrocytes"

### 0001379_01_TH.jpg

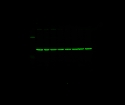

### 0001380_01_TH.jpg

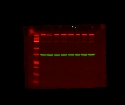

### 0001381_01_TH.jpg

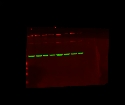

### 0001392_01_TH.jpg

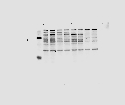

### 0001393_01_TH.jpg

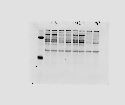

### 0001394_01_TH.jpg

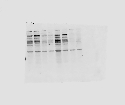

### 0001940_01_TH.jpg

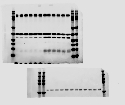

### 0001941_01_TH.jpg

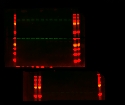

### 0001953_01_TH.jpg

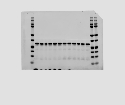

### 0001954_01_TH.jpg

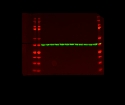

### 0001955_01_TH.jpg

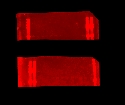

### 0002120_01_TH.jpg

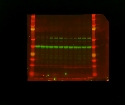

### 0002122_01_TH.jpg

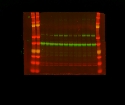

### 0002127_01_TH.jpg

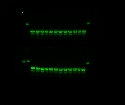

### 0002191_01_TH.jpg

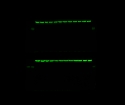

### Exp247b_set2_SyproRuby.raw16.tif

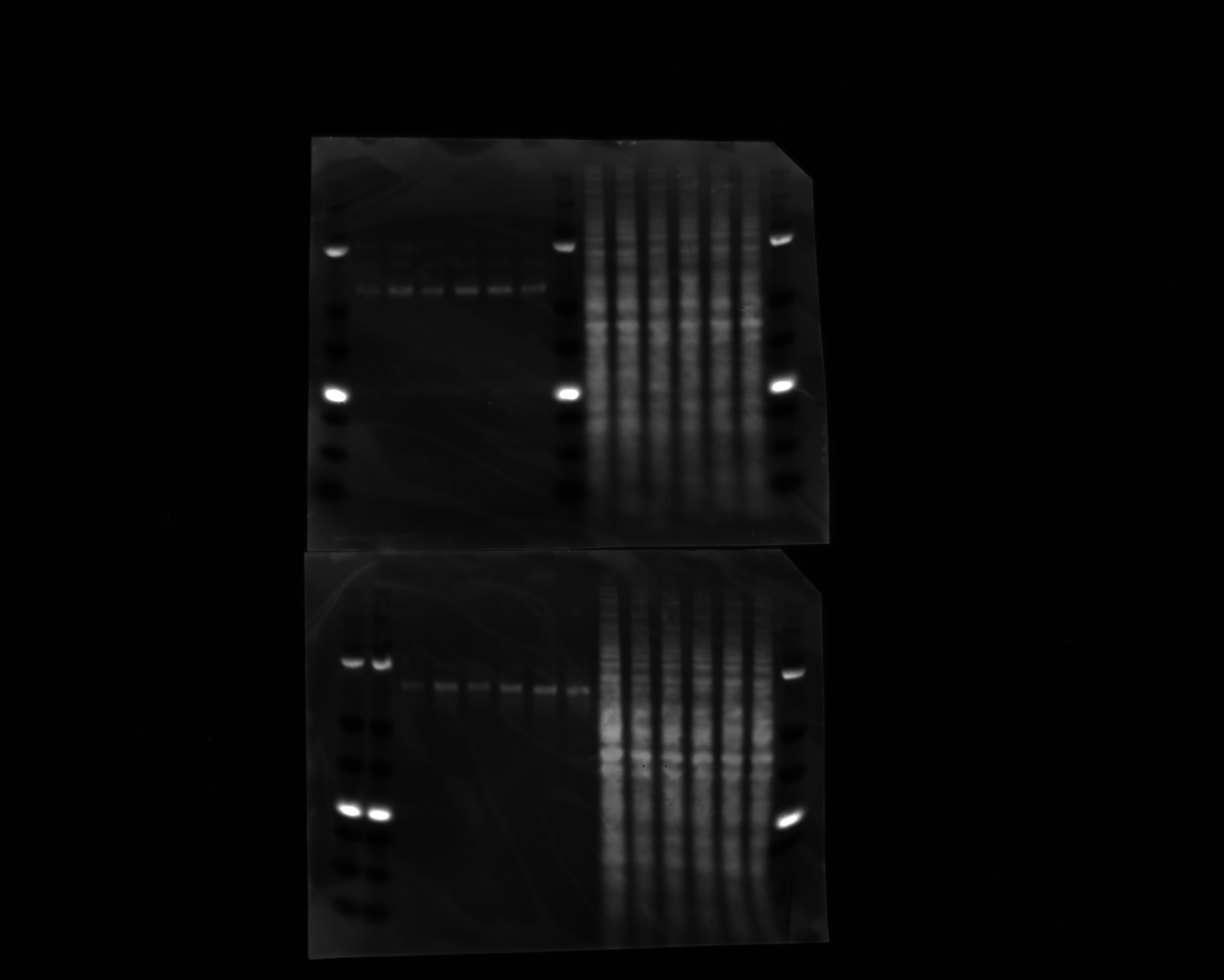

### Exp247c_set1_SyproRuby.raw16.tif

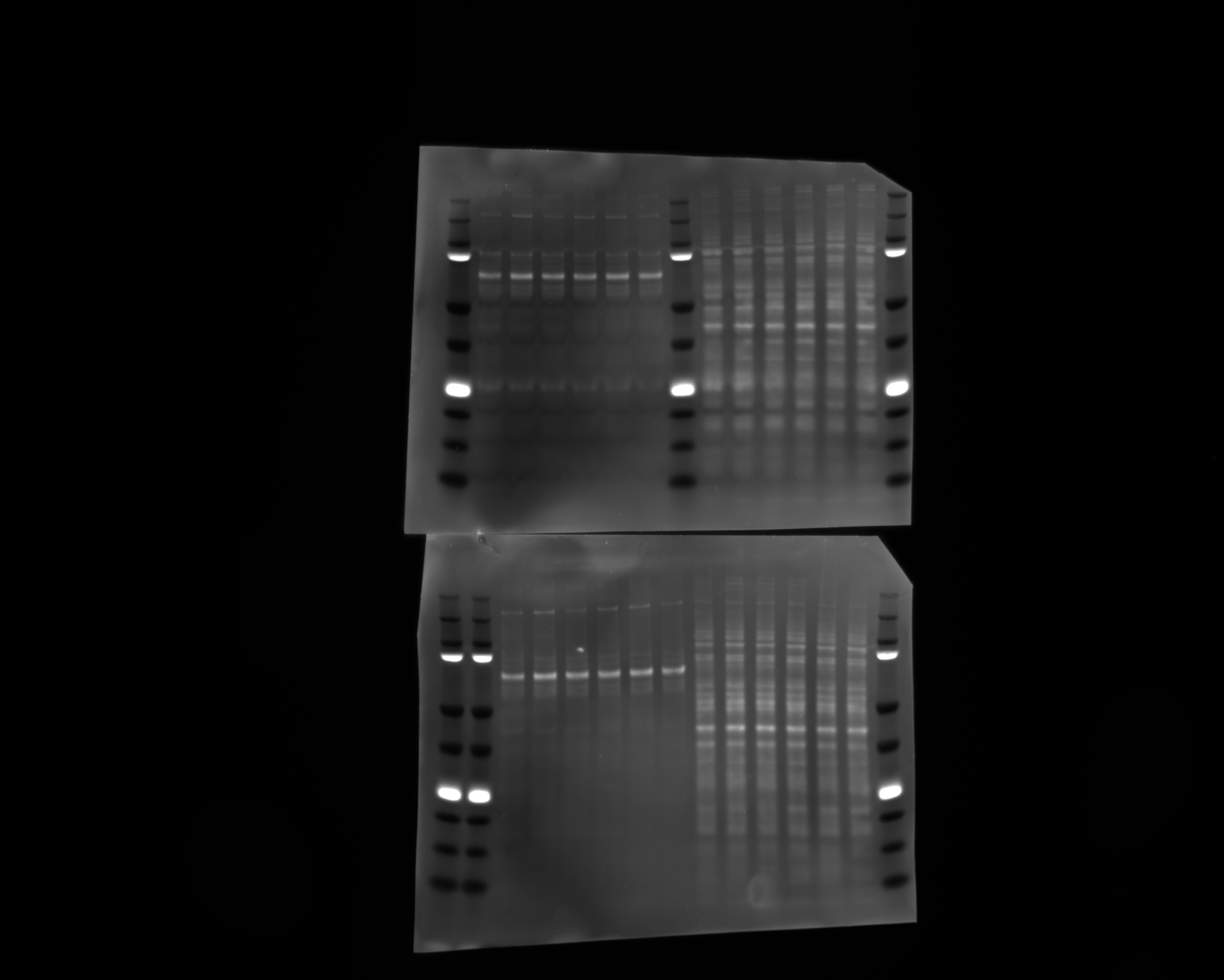

### movie2_quantification.pdf

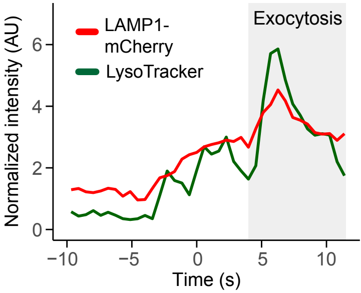

### movie2_selectedFrames.pdf

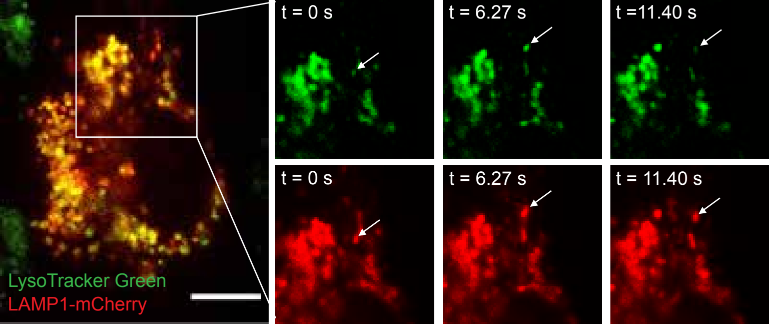

### syproRUBY.raw16.tif

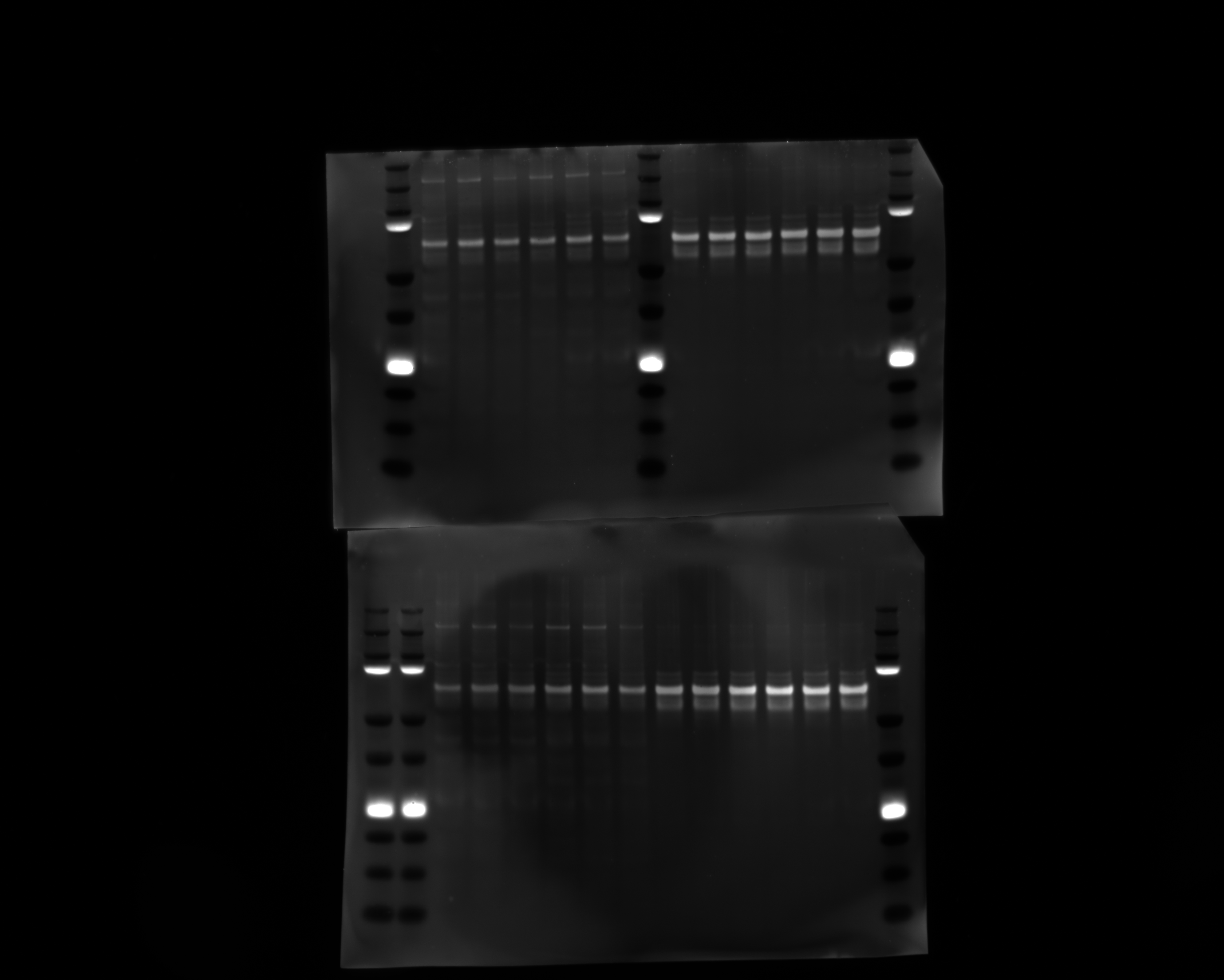
